# MHC-I diversity enables rapid adaptation during viral pandemic in wild rabbit populations

**DOI:** 10.1101/2025.10.30.685367

**Authors:** Yexin Zhang, Jonathan P. Day, Marina Lirintzi, Jiayi Ji, Clive Tregaskes, Miguel Carneiro, Joel M. Alves, Tanja Strive, Jim Kaufman, Francis M. Jiggins

**Affiliations:** Department of Genetics, University of Cambridge, Cambridge CB2 3EH, United Kingdom; Department of Genetics, Evolution and Environment, University College London, London WC1E 6BT, United Kingdom; CIBIO, Centro de Investigação em Biodiversidade e Recursos Genéticos, InBIO Laboratório Associado, Campus de Vairão, Universidade do Porto, Vairão, Portugal; BIOPOLIS Program in Genomics, Biodiversity and Land Planning, CIBIO, Campus de Vairão, Vairão, Portugal; Palaeogenomics and Bio-Archaeology Research Network, School of Archaeology, University of Oxford, Oxford, OX1 3QY, United Kingdom; Commonwealth Scientific and Industrial Research Organisation, Health and Biosecurity, Canberra, Australia; Institute for Immunology and Infection Research, School of Biological Sciences, University of Edinburgh, Edinburgh, United Kingdom; Department of Immunobiology, Yale University, New Haven, Connecticut 06510, USA

## Abstract

Emerging diseases can have devastating consequences for wild species, with long-term effects depending on the ability of the host to evolve resistance. Here, we show that major histocompatibility complex (MHC) genes provided standing genetic variation that enabled rabbits to mount a rapid evolutionary response to the myxoma virus pandemic that began in the 1950s. Using historical and modern specimens starting in 1865 and spanning the pandemic, we found strong parallel shifts in MHC-I allele frequencies across Australia, Britain and France, alongside population-specific signals. These evolutionary shifts appear to have altered antigen presentation to T cells. Our results provide evidence that MHC-I is under strong selection in natural populations during a pandemic, and that the high polymorphism of MHC may have contributed to the evolutionary rescue of these populations.

## Introduction

The emergence of a new infectious disease when a novel pathogen arrives in a population can potentially lead to population collapse or even extinction. Its impact may be especially severe because the host has not evolved resistance or tolerance. Therefore, the long-term effects will depend on the population’s ability to adapt, which can rescue it from decline or extinction^1^. Adaptation may proceed primarily from standing genetic variation already present in the population, from novel beneficial mutations that occur after pathogen arrival, or from a combination of both^2^. The relative importance of these mechanisms remains an enduring question in evolutionary biology, with important implications for understanding how populations persist in the face of emerging infectious diseases as well as ecological change.

A potential source of genetic variation underlying resistance or tolerance to infections is the Major Histocompatibility Complex (MHC), which typically represents the most polymorphic gene region in vertebrate genomes, encoding cell surface molecules that present pathogen-derived peptides to T cells and thereby initiate adaptive immune responses^3^. Classical MHC class I (MHC-I) genes are expressed on all nucleated cells and primarily present intracellularly derived peptides, making them essential for detecting viral infections^4^. Different alleles of MHC-I encode molecules that present different pathogen-derived peptides, which can lead to differences in susceptibility to infection^5^.

The extraordinary diversity maintained within MHC genes results from balancing selection maintaining multiple alleles through mechanisms including heterozygote advantage, negative frequency-dependent selection, and fluctuating selection driven by temporal and spatial variation in pathogen pressure^6,7^. Evidence for these processes comes from HIV in humans, where disease progression is slower in individuals that are heterozygous for MHC-I (HLA-I)^8^ or who carry rare alleles^9,10^. In sticklebacks, locally common MHC alleles tend to be protective against local parasites^7^. However, much evidence that MHC underpins adaptation during pathogen outbreaks remains indirect^11–13^. Demonstrating real-time selection, though, requires temporal comparisons of allele frequencies before and after pathogen emergence. Such temporal sampling is rarely available for wild populations experiencing natural disease outbreaks. Even when temporal data are available, the unknown identity of the selective agents often precludes definitive attribution to pathogen-driven selection^14^.

The European rabbit-myxoma virus system provides an exceptional opportunity to study MHC evolution during a documented viral pandemic. Myxoma virus (MYXV) was deliberately introduced to control rabbit populations in Australia (1950), France (1952), and Britain (1953), causing catastrophic mortality with case fatality rates exceeding 99%^15–17^. This created a natural experiment replicated across three populations, with extensive historical records and museum specimens providing genetic material from populations before virus exposure. Previous genome-wide studies identified evolutionary changes within the MHC region following MYXV introduction^18^. However, the specific genes under selection in this region and their functional significance remained unclear due to missing data and poor annotation.

Here, we study how MHC-I alleles affected rabbit fitness following the release of MYXV, asking whether natural selection produced a directional shift favouring individual alleles or instead promoted a broader shift in the frequency of multiple alleles consistent with balancing or frequency-dependent selection. Our findings reveal that two classical MHC-I genes were targets of intense selection, with specific haplotypes showing large parallel frequency changes across populations. These results provide unprecedented insight into how immune gene diversity responds to novel pathogen challenges and demonstrate the critical importance of standing genetic variation in enabling rapid evolutionary responses to emerging diseases.

## Results

### Reconstruction of rabbit MHC region reveals two clusters of MHC-I genes

The emergence of MYXV was accompanied by the rapid evolution of several immunity genes across the rabbit genome^18^. Although we previously reported changes in non-coding SNP allele frequencies within the MHC region^18^, poor annotation of this complex gene-dense locus and limited population-genetic data mean that the specific targets of natural selection within the MHC remain unknown. To address this, we designed hybridisation probes to capture rabbit MHC transcripts, enabling us to generate full-length transcript sequences using PacBio HiFi long-read sequencing from 68 rabbit samples (data file S1), including wild rabbits from Australia and Britain, domestic breeds and a rabbit kidney cell line (RK13).

To annotate MHC-I genes, we mapped these transcript sequences to the rabbit reference genome OryCun 2.0. This revealed four previously unannotated MHC-I genes and errors in most of the existing gene annotations. We therefore manually annotated these genes, revealing nine MHC-I genes on chromosome 12, which we named *Orcu-U1* to *Orcu-U9* (Fig. 1A, table S1, data file S2 and data file S3). Phylogenetic analysis confirms that these nine genes are more closely related with human MHC-I genes than to other Class-I-like genes (Fig. 1B and data file S4). The first four are tightly clustered within a 100 kb region, while the subsequent five are grouped within a 300 kb region, separated by an interval of approximately 2 Mb (Fig. 1A). This two-cluster organisation reflects a translocation. Humans likely represent the ancestral mammalian state, consisting of a continuous MHC-I region (including *Extend MHC1, Alpha, FW1, Kappa, FW2* and *Beta*), followed by MHC-III and MHC-II clusters^19^. In rabbits, a portion of the MHC-I region (partial *FW2,* and *Beta*) was translocated along with the MHC-III cluster to a position in the middle of the Extended MHC-I region (Fig. 1C and fig. S1). Unlike the second cluster of MHC-I genes (*Orcu-U5* to *Orcu-U9*; fig. S2b), the first region containing *Orcu-U1* to *Orcu-U4* showed a high degree of repetition, reflecting the origin of these genes from a series of duplications (Fig. 1D and fig. S2a). A large, inverted repeat marks a duplication event that gave rise to *Orcu-U1* and *Orcu-U2*, and a shorter duplication within this region gave rise to *Orcu-U3* (Fig. 1D). This reconstruction of the MHC-I locus provides the essential genomic framework to for investigating the targets of natural selection within this critical immune region.

**Fig. 1.**
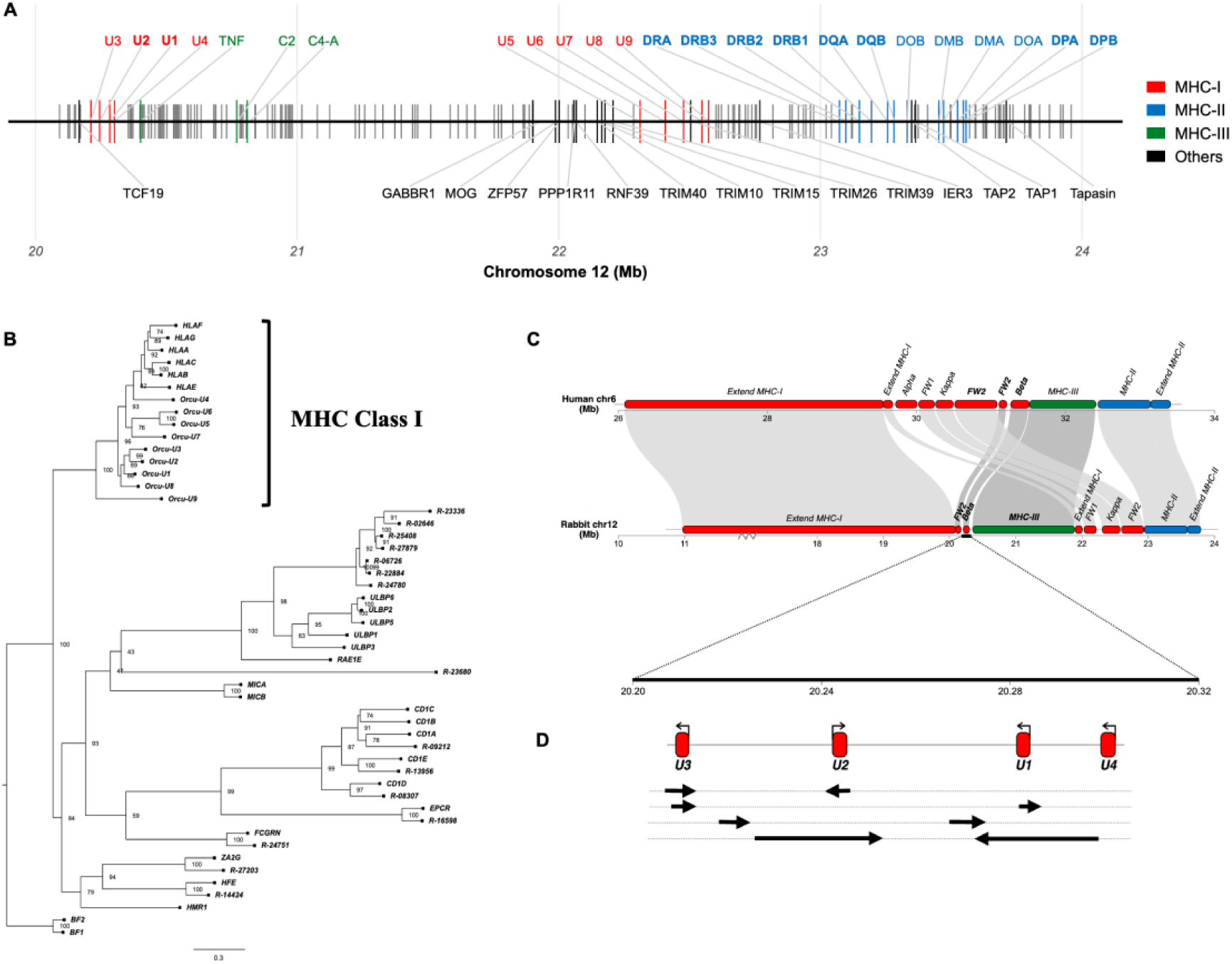
MHC-I region of European rabbits. (**A**) Physical positions of nine MHC-I genes on chromosome 12 of the rabbit genome OryCun 2.0. (**B**) Maximum likelihood phylogeny (GTR20 model) of MHC-I classical and non-classical, and MHC-I-like protein sequences of humans and rabbits. Branch support is indicated as percentage of 1,000 bootstrap replicates. The genes with names starting as ‘*Orcu’* are annotated here using transcript sequencing data, while those starting by ‘R’ are annotations from the *Ensembl* database. *HLA-A, HLA-B, HLA-C* are human classical MHC genes and *HLA-E, HLA-F, HLA-G* are non-classical. Other proteins share the basic MHC-I structure but function beyond classical peptide presentation to T cells. Chicken MHC-I genes (*BF1* and *BF2*) were used as outgroups to root the tree. (**C)** Chromosomal rearrangement between human (human assembly GRCh38.p13 chr6; top) and rabbit MHC (rabbit assembly OryCun2.0 chr12; bottom). (**D**) A zoomed-in view of the MHC-I genes located in the *Beta* region of the rabbit genome. The red segments mark the genomic positions of the genes, while the curved arrows indicate their transcriptional direction. In the underlying tracks, four pairs of arrows highlight the tandem duplication events that occurred within this region.

### Natural selection maintains extreme polymorphism in two classical MHC-I molecules

In other species, only a subset of MHC-I genes encodes classical molecules responsible for presenting a diverse range of pathogen peptides to T cells^20,21^. These genes have conserved sequence motifs required for functions including peptide binding and T cell recognition. To identify these, we aligned the human and rabbit proteins (Fig S3). The signal peptide of *Orcu*-*U3* and *Orcu*-*U8* terminates in TRE or TRT, violating signal peptidase specificity rules, which require small residues (A/G/S/C/P) at the -1 position and helix-compatible residues (A/G/S/C/T/V/I/L/P) at -3 for efficient cleavage^22,23^, suggesting defective processing (Fig S3). Furthermore*, Orcu-U3* lacks the N-X-S/T glycosylation motif within the α1 domain, while *Orcu*-*U9* gene has a run of extra residues after this motif^24^. Such structural deviations would disrupt UGT1/TAPBPR-mediated quality control—a critical checkpoint for classical MHC-I folding and peptide loading^24,25^.

Additional non-classical hallmarks include: 4-residue deletions in the α2 domain of *Orcu-U5/U6*, an atypical α3-domain loop in *Orcu-U7*, and truncated cytoplasmic tails in *Orcu-U4/U9* possibly resembling the immunosuppressive HLA-G^26^. Notably, *Orcu*-*U5* and *Orcu*-*U6* also lack the conserved tyrosine-based YXXØ motif which functions as an endocytic trafficking signal that is critical for MHC class I internalisation and cross-presentation of exogenous antigens in dendritic cells^27,28^. In contrast, *Orcu*-*U1* and *Orcu*-*U2* retain all canonical structural features characteristic of classical MHC-I molecules, including the invariant residues that coordinate N- and C-termini of the bound peptide (Y7, Y59, Y159, Y171 and Y84, T143, K146, W144; fig. S3)^29^, suggesting a primary role in conventional antigen presentation and competence for peptide presentation to T cells. Because classical MHC-I genes are also highly expressed across all nucleated cells, we next used RNA sequencing data to investigate expression levels (Fig. 2A and table S2). This revealed that *Orcu*-*U1* and *Orcu*-*U2* were both highly expressed across multiple tissues, consistent with a role as classical MHC-I genes. The mean expression of *Orcu*-*U1* was 4 times greater than *Orcu*-*U2. Orcu*-*U8* was also highly expressed across tissues, while other genes had lower and/or more tissue-specific expression.

**Fig. 2.**
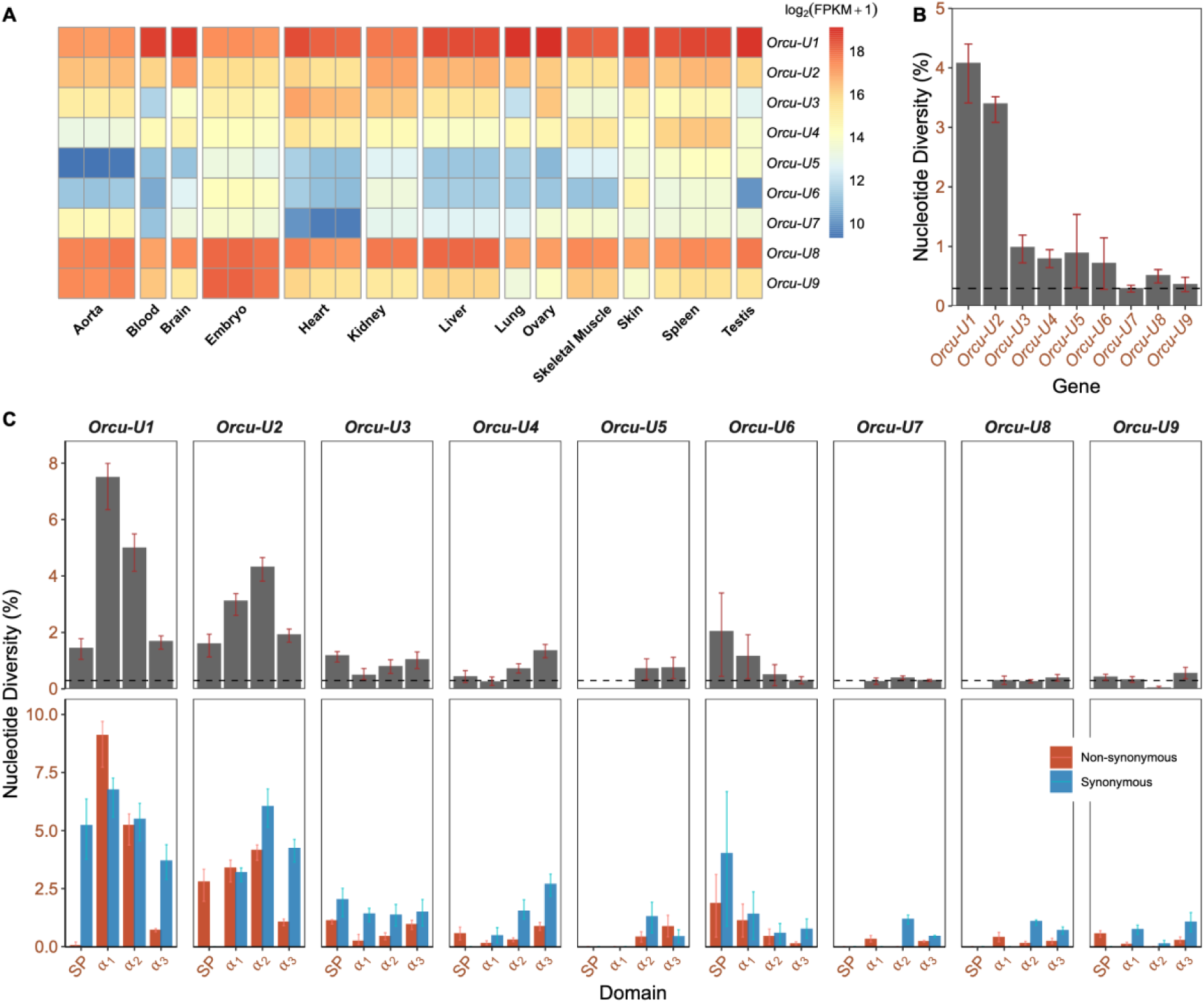
Genetic diversity and expression of MHC-I genes. **(A)** Gene expression across tissues and MHC-I genes estimated from RNA sequencing data. **(B)** Nucleotide diversity of 9 MHC-I genes in Britain. **(C)** Nucleotide diversity, non-synonymous and synonymous nucleotide diversity of different exons encoding different domains in Britain. The domains are the signal peptide (SP), the α₁ and α₂ domains which together form the peptide-binding groove, and the transmembrane-proximal α₃ domain. The top panel shows total diversity, while the bottom panel distinguishes between non-synonymous and synonymous diversity. The dashed horizontal line on panels **(B)** and the top row of **(C)** indicates the mean genome-wide nucleotide diversity. Confidence intervals in **(B)** and **(C)** correspond to the 0.025 and 0.975 quantiles of 1,000 bootstrap replicates estimates obtained by resampling alleles with replacement.

The third hallmark of classical MHC-I genes is a high degree of polymorphism, especially in the domains that bind pathogen peptides. Using our long-read transcript sequences, we identified 12, 9, 13, 10, 5, 5, 9, 9, and 7 nucleotide alleles across *Orcu*-*U1* to *Orcu*-*U9*, respectively. The genes *Orcu*-*U1* and *Orcu*-*U2* have very high nucleotide diversity, exceeding that of the other MHC-I genes (Fig. 2B, fig. S5 and fig. S6a) and sequences elsewhere in the rabbit genome (Fig. 2B and fig. S5)^30^. Furthermore, the most variable regions in these genes are exons 2 and 3, which encode the α1 and α2 domains involved in binding and presenting peptides to T cells (Fig. 2C and fig. S6b). Adaptive pressure exerted by pathogens on classical MHC genes drives an elevated level of non-synonymous polymorphism at the codons coding for residues contacting pathogen-derived peptides^31^. Consistent with this, *Orcu*-*U1* and *Orcu*-*U2* display substantially elevated non-synonymous diversity within their peptide-binding α1 and α2 domains when contrasted with the other MHC-I genes (Fig. 2C and S6b). These domains also have elevated synonymous nucleotide diversity, as expected if the alleles are ancient and have been maintained by balancing selection (Fig. 2C and fig. S6b). The high diversity observed in *Orcu*-*U1* and *Orcu*-*U2* supports our conclusion that these are classical MHC-I genes and indicates that they harbour functional variation that affects pathogen recognition.

### Natural selection drove parallel and population-specific changes in MHC-I during the MYXV pandemic

To understand how selection has acted on MHC-I over the course of the MYXV pandemic, we compared allele frequencies between pre-pandemic museum specimens and contemporary samples. However, this presented two challenges: First, our sequencing strategy for full-length MHC transcripts requires high molecular weight RNA from fresh tissue, which is not available for historical material. Second, phylogenetic trees constructed from individual exons revealed that alleles from different genes often intermingle, with their positions varying by exon (fig. S7). This suggests that sections of DNA are exchanged by gene conversion between alleles of the different genes^32^. As a result, it is not possible to reliably assign short sequence reads to specific genes using conventional mapping approaches.

To overcome these issues in modern rabbits (data file S1), we amplified three overlapping regions covering exons 1-3 using PCR primers that were conserved across *Orcu*-*U1* and *Orcu*-*U2*. The amplicons were then sequenced using Illumina sequencing, and we developed a pipeline to assemble the phased sequences of different alleles (see methods). Within the signal peptide (exon 1), there are fixed differences between *Orcu*-*U1* and *Orcu*-*U2,* so this was used to assign alleles to a specific gene, while exons 2 and 3 contain most of the sequence variation. We confirmed the accuracy of this method by including the individuals for which we had generated complete transcript sequences, and in all cases the sequences and genotypes of *Orcu*-*U1* and *Orcu*-*U2* were identical between the two approaches. Combined with the transcript analysis above, this allowed us to identify 29 alleles of *Orcu*-*U1* and 17 alleles of *Orcu*-*U2* from 161 rabbits. Thus, 104 nucleotide alleles were identified across all 9 MHC-I paralogs (table S1, data file S3 and data file S5).

Due to the high level of DNA degradation of the museum samples, which prevented a standard PCR amplification we designed new hybridisation probes that incorporated the allelic diversity found in modern samples and used these to capture the MHC-I from 95 historical Illumina sequencing libraries (data file S1). As gene conversion prevents us from reliably mapping the reads from historical specimens to specific genes, we implemented a modified methodology often used in human HLA typing^33^. We mapped the reads from each rabbit onto all known MHC-I alleles across all eight MHC-I genes from *Orcu-U1* to *Orcu-U8* and selected the 1-2 alleles across each locus that maximises the total number of mapped reads. To optimise and validate this approach, we simulated sequence reads with the same patterns of fragmentation and post-mortem mutation that we observed in our museum dataset. Simulated reads were generated for all eight MHC-I genes and across the set of genotypes of that we observed in the modern individuals. Based on these simulations we required a 97% match between a read and the target sequence to map the read and selected a penalty for calling heterozygotes that effectively mitigated a bias toward heterozygosity (beta=0.010). We found that ∼97% of alleles were correctly called, provided there was a minimum coverage of 3X (fig. S8). We applied this approach to 79 rabbits from our historical collection with >3X coverage, collected between 1865 and1956 in France (*n*=30), Britain (*n*=32) and Australia (*n*=17) (data file S3).

Next, we compared allele frequencies between modern and museum specimens from similar geographical locations in Britain, France and Australia. Our previous work has scanned the rabbit genome for sites where natural selection had driven changes in allele frequency^18^. As sequences from the MHC-I coding regions were either missing or unreliable in this dataset (see above), we added our sequences from exon 2 and 3 of *Orcu-U1* and *Orcu-U2* from modern and historical individuals to the analysis. We modelled allele frequency changes through time using an approach that accounts for historical patterns of admixture and identified significant changes in SNP allele frequencies that were driven by natural selection using a likelihood ratio test^18,34^. Using a model which allows strength of selection in the three populations to vary independently, we found six SNPs in the coding region of *Orcu-U1* and *Orcu-U2* where natural selection had driven changes in allele frequency that were significant at the genome-wide level (Fig. 3A). A model assuming the selection coefficient was the same across three populations showed similar pattern (fig. S9). There was no evidence of natural selection acting on the other seven MHC-I paralogs (Fig. 3A, fig. S9). Therefore, natural selection has acted on classical MHC-I genes during the MYXV pandemic.

**Fig. 3.**
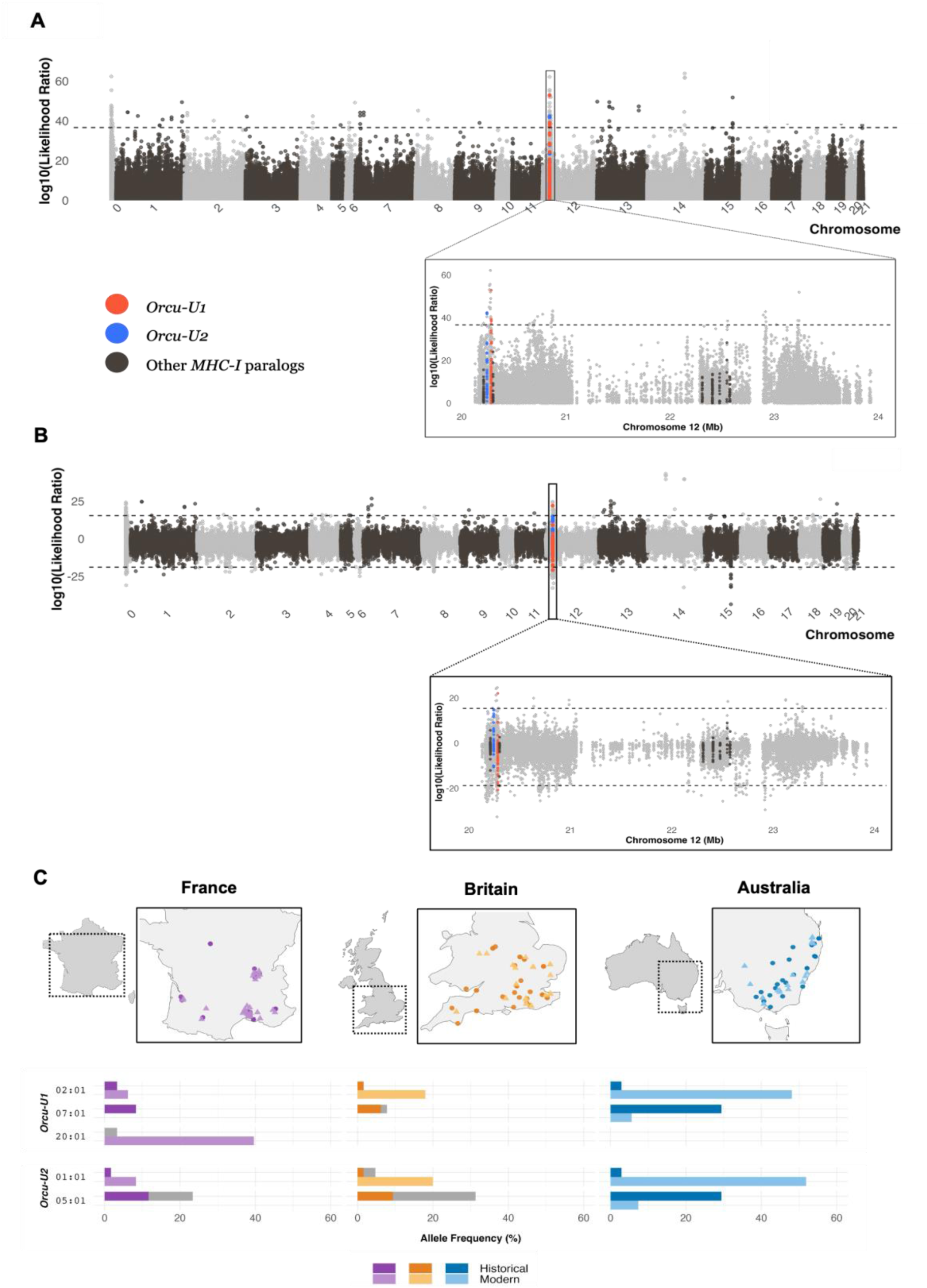
Parallel and population-specific changes in allele frequency across three countries. **(A)** Selection scan based on allele frequency changes after the introduction of myxomatosis (the strength of selection in each population is allowed to vary independently). (**B**) Selection scan testing whether selection has acted in all three populations (positive values) or just one population (negative values). (**A**) and (**B**) The plot shows the genome-wide analysis, and the inset the MHC-I region, with the red and blue dots showing variants within the exon region of *Orcu-U1* and *Orcu-U2*, and the brown dots representing the variants within the exon regions of the other seven MHC-I paralogs. The y axis shows the likelihood ratio statistic of each model. Dashed black lines indicate the genome-wide 95% significance threshold from permuting sample collection dates within each country 1,000 times. **(C)** The maps show historical (circles; 1856-1956) and modern (triangles; 2002-2013) sampling locations in France, Britain and Australia. The bottom panel demonstrates the allele frequency changes before and after myxomatosis within three populations. The allele frequencies for each allele in gene *Orcu-U1* and *Orcu-U2*, in modern and historical samples from France, Australia, and Britain. Only alleles with significant changes in frequency shown. Alleles from historical specimens represented with stacked grey bars resemble but are not identical to that sequence in our database.

Britain, Australia and France have all experienced intense selection by myxomatosis over this time, so parallel changes in allele frequency across populations might be expected if this disease underlies the selection acting on *Orcu-U1* and *Orcu-U2.* We therefore tested whether a model of parallel selection or population-specific selection better explained the genetic data. We found a variant in *Orcu*-*U1* with parallel changes across populations (Fig. 3B; positive values), and another variant from the same gene having population-specific changes (Fig. 3B; negative values).

The properties of an MHC molecule depend on the entire sequence rather than isolated SNPs, so we compared *Orcu-U1* and *Orcu-U2* allele frequencies before and after myxomatosis (Fig. 3C and fig. S10). Overall, there are significant changes of allele frequency between historical and modern populations within each gene and population (Fisher’s Exact Test; *Orcu-U1: p* = 7.61×10^-10^, *p* = 0.024, *p* = 1.30×10^-6^; *Orcu-U2*: *p* = 0.002, *p* = 5.00×10^-4^, *p* = 9.72×10^-7^, for France, Britain, and Australia respectively). Notably, there are two alleles, *Orcu*-*U1\**02:01 and *Orcu-U2\**01:01, showing parallel increases in frequency in all three populations. The changes in frequency were large, with *Orcu-U1\**02:01 increasing from 2.9% to 48.1% in Australia. This allele was present at a frequency of 3.3%, 1.6%, and 2.9% in France, Britain and Australia before the myxomatosis outbreak, so the high polymorphism of MHC-I genes may have provided standing genetic variation that could be selected by myxoma virus, contributing to the rapid evolution of resistance to myxomatosis after the outbreak.

Conversely, *Orcu-U1\**07:01 and *Orcu-U2\**05:01 exhibited a parallel decrease in frequency across all three populations. Strikingly, these two alleles were relatively common in all populations before the introduction of MYXV but have disappeared entirely from modern samples in France and Britain. In Australia, while still present, their frequencies have dramatically decreased from 29.4% to 5.6%, and 29.4% to 7.4%, respectively. The sharp decline of these previously common alleles suggests they may have conferred susceptibility to the myxoma virus and became disadvantageous under the intense selection pressure exerted by myxomatosis.

In France, the allele *Orcu*-*U1\**20:01 was absent from the historical sample but rose to 39.6% frequency in the modern dataset (Fig. 3C). This allele was not detected in any of the samples from Australia and Britain, suggesting it was likely lost when rabbits were introduced in Britain during the medieval period^35^ and from there transported to Australia in the 19^th^ century^36^. Consequently, this population-specific adaptation likely resulted not from distinct selection pressures but rather from differences in the MHC-I alleles available for selection to act upon. The peptide-binding domain of *Orcu-U1*\*20:01 is genetically distant from *Orcu-U1*\*02:01—the allele that increased in frequency across all three populations—suggesting that these alleles are unlikely to be functionally equivalent (fig. S11). The large increase in the frequency of *Orcu-U1*\*20:01 in France, may explain why the *Orcu-U1*\*02:01 and *Orcu-U2*\*01:01 haplotype had a smaller increase in frequency in here than the other populations (Fig. 3C) and showed a strong signal in a population-specific selection scan (among the top 0.003% of variants in the genome, fig. S12).

As the two classical MHC-I genes are adjacent in the genome, there is the potential to generate an MHC-I ‘supergene’ if alleles of the two genes are tightly associated, potentially increasing the strength of disease associations. We found strong linkage disequilibrium between alleles of *Orcu-U1* and *Orcu-U2,* including those that had experienced the greatest changes in frequency (fig. S13). Using statistical phasing^37^, we found that the alleles that experience large and parallel changes in allele frequency were overwhelmingly found on the same haplotype—we estimate that 97.5% of *Orcu*-*U1\**02:01 alleles were on a haplotype that carried *Orcu-U2*\*01:01, and 90.8% of *Orcu-U2\**01:01 alleles were on a haplotype that carried *Orcu*-*U1*\*02:01 (data file S6). Similarly, the alleles that decreased in frequency were also found on a single haplotype (*Orcu-U1\**07:01 and *Orcu-U2\**05:01; data file S6). Therefore, the strongest signal of selection reflects changes in the frequency of two haplotypes, each of which encompasses both classical MHC-I genes.

### Selection was intense and acted upon MHC-I before the emergence of RHDV

In 1984, another lethal disease was identified in rabbits, caused by the positive-sense RNA virus rabbit haemorrhagic disease virus (RHDV; *Caliciviridae*). It was first detected in domestic rabbits in China, and it has a similarly high case fatality rate to MYXV^38,39^. The virus was detected in continental Europe, Britain, and Australia, in 1986, 1992, and 1995, respectively, and in all these locations it caused considerable mortality^39–41^. To evaluate the role that RHDV has been playing in selection on different MHC-I alleles through time, we used amplicon sequencing to genotype 70 rabbit samples collected before or shortly after RHDV spread in Britain and Australia populations (1985 to 1996; data file S1).

Focusing on the haplotypes with the greatest change in frequency, we used a Bayesian approach^42^ to test whether selection was acting before the emergence of RHDV. To do this, we estimated separate selection coefficients from the time when MYXV was released to the appearance of RHDV, and from the appearance of RHDV to the present day. We found a significant signal of positive selection acting on the haplotype encompassing two alleles *Orcu-U1\**02:01 and *Orcu-U2\**01:01 from before the release of RHDV in both Britain and Australia. Furthermore, rabbits carrying this haplotype enjoyed a considerable advantage, with a per-year selection coefficient of 0.06 and 0.08 in Britain and Australia, respectively (Fig. 4A and data file S7). There was no significant change in the selection coefficient after the release of RHDV. The haplotype with the largest decrease in frequency (*Orcu-U1\**07:01*, Orcu-U2*\*05:01) also showed evidence of selection before RHDV in both populations, with a selection coefficient -0.1 in Australia, which is the population where we have the most data to estimate this parameter. These results therefore indicate that myxoma virus was likely the principal selective agent driving these frequency changes of these two haplotypes.

**Fig. 4.**
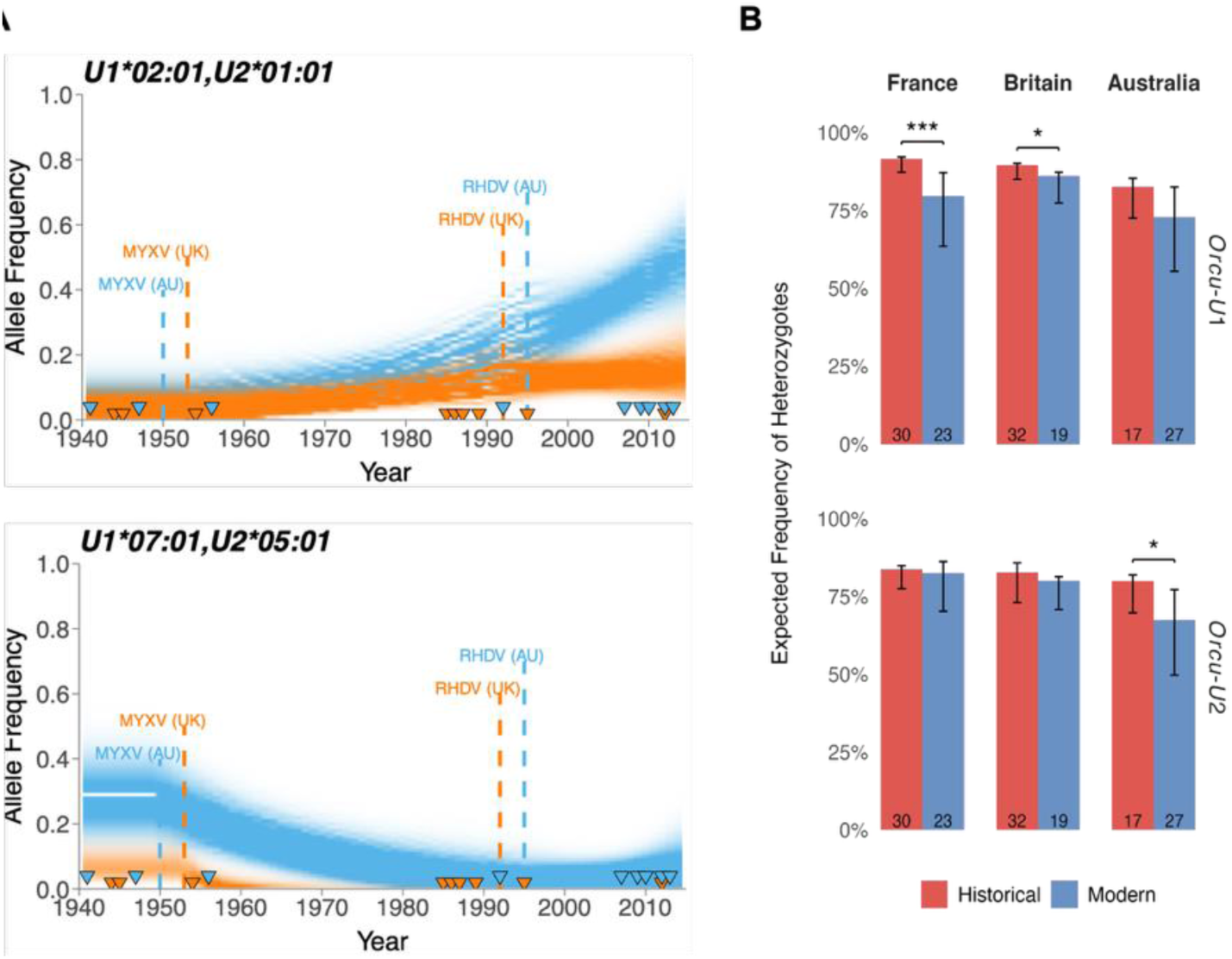
Changes in selection strength and heterozygosity across populations and time periods. (**A**) Posterior distribution of estimated haplotype frequencies through time for the positively and negatively selected haplotypes of *Orcu-U1* and *Orcu-U2* inferred from the Bayesian selection analyses. Shaded areas show estimated selection strength with 95% credible intervals for UK (orange) and Australia (blue), assuming effects on fitness are additive. Dashed lines show the dates of the first observations of MYXV and RHDV. Triangles at the bottom represent the years of samples used in this analysis. (**B**) Bars represent expected heterozygosity of *Orcu-U1* and *Orcu-U2* in historical and modern samples calculated from observed allele frequencies. Sample sizes are indicated at the base of each bar. Error bars show 95% confidence intervals estimated via 1,000 bootstrap replicates by resampling samples with replacement. Statistical significance of differences between historical and modern groups was assessed using 10,000 permutation tests where time period labels (Historical vs Modern) were randomly reassigned to individuals to generate a null distribution (\**p<0.05*, \*\*\**p* < 0.001).

Individuals who are heterozygous for MHC-I alleles can present a wider array of peptides and may be more resistant to infection^8^. However, we can reject this as the main selection pressure driving MHC-I evolution during the MYXV pandemic because the changes in allele frequency we observed result in significant decreases in heterozygosity across multiple populations (Fig. 4B). This reduction in genetic heterozygosity suggests that selection acted directionally to increase the frequency of specific beneficial alleles, rather than maintaining balanced polymorphism to maximise the overall probability of heterozygosity. However, natural selection may instead favour individuals with more divergent alleles because the sequence divergence between alleles predicts how similar the repertoire of peptides presented is^43–45^. In Australia, evolution increased the expected divergence between alleles within individual rabbits, with a non-significant trend in this direction in the other two populations (fig. S14a). This was the result of the positively selected allele *Orcu-U2\**01:01 being the most divergent allele in all three populations, while *Orcu-U1\**02:01 also had high divergence (fig. S14b). Therefore, selection favoured a haplotype carrying a divergent allele.

## Discussion

Our study provides direct evidence for rapid, parallel adaptive evolution of classical MHC-I genes during the myxoma virus pandemic in European rabbits across three populations and two continents. By integrating historical museum specimens and modern samples spanning over 150 years, we demonstrate that standing genetic variation in MHC genes enabled parallel evolutionary responses to a novel pathogen across geographically isolated populations, offering unprecedented insights into real-time evolution during a global disease outbreak. Importantly, we demonstrate direct evidence of strong pathogen-driven selection on MHC in a wild population.

A striking finding is the parallel genetic changes MHC-I across Australia, Britain, and France, with all three showing remarkably similar trajectories of allele frequency change following myxoma virus introduction. The most strongly selected alleles existed at frequencies below 4% in pre-pandemic populations, but increased dramatically to become dominant in Britain and Australia. This convergent response suggests that the selective pressure was the myxoma virus, as all three populations experienced extreme selection from the virus at this time and all responded by evolving strongly increased resistance^46–48^. Furthermore, despite a large family of MHC genes being identified and annotated, natural selection was focused on the two classical MHC-I genes, indicating that these changes shaped antigen presentation to T cells.

While parallel evolution dominated the response across continents, one of the largest changes was the population-specific rise of *Orcu-U1*\*20:01 to high frequency in France. This allele was absent from our samples in Australia and Britain. This suggests that the allele may have been lost when rabbits were introduced to Britain from France ∼800 years ago and likely represents an alternative solution to the same selective challenge, illustrating how historical contingencies can shape adaptive trajectories.

The dramatic population crashes observed during the myxoma pandemic serve as a stark reminder of how emerging diseases can threaten species survival, even in abundant and widespread populations. When novel pathogens enter a species or population, they often have especially severe impacts because the host population has not yet evolved resistance or tolerance. The long-term outcome therefore depends on the host’s ability to adapt, which may allow populations to recover^1^. This is exemplified by rabbit populations, where long-term monitoring data from Australia documented repeated pandemic-driven collapses following introductions of new pathogens, pathogen variants and disease vectors^49^. However, populations consistently recover after ∼10-15 years. These patterns suggest that host resistance evolution, rather than pathogen attenuation, may drive population recovery^50^. This adaptive potential, in turn, relies on both the amount and nature of genetic variation in susceptibility to the infection. Our results suggest that the standing genetic variation maintained at MHC loci provides a mechanistic basis for this evolutionary rescue, allowing populations to rapidly adapt to successive pathogen challenges and persist despite repeated disease outbreaks that would otherwise lead to extinction.

MHC may play a crucial role in adaptation to novel pathogens by maintaining high levels of standing genetic variation. Standing variation allows for rapid evolutionary responses, as beneficial alleles already exist within the population and are at higher initial frequencies than new mutations. In the case of MYXV, which spread over thousands of kilometres within months and had an initial fatality rate of 99.8%^17^, the rapid host evolutionary response would likely have been impossible if populations had to wait for beneficial mutations to arise de novo. The MHC alleles that increased in frequency during the pandemic were pre-existing variants. From a conservation perspective, these findings highlight the importance of preserving MHC genetic diversity in wild populations. Conservation strategies should therefore focus not only on maintaining population sizes but also on safeguarding genetic diversity—particularly in MHC may provide resilience against future emerging diseases. That said, our results suggest that only extreme and prolonged reductions in population size may significantly erode this diversity, as most Australian rabbits today are descended from a small number of individuals^36^.

Natural selection has influenced the frequencies of haplotypes containing alleles from both *Orcu-U1* and *Orcu-U2*. The most strongly selected alleles of the two genes are overwhelmingly found together on the same chromosome, creating a haplotype that can be viewed as a single MHC-I supergene. This may strengthen the effects of selection, and the observed association could itself result from fitness epistasis if it is advantageous to carry both alleles simultaneously. However, due to the limited number of recombinants in our samples, we unable to test this hypothesis, and selection may be acting primarily on one gene, with the associated allele at the other locus increasing in frequency through genetic hitchhiking.

The dramatic shifts in MHC allele frequencies following myxomatosis may have disrupted previous evolutionary equilibria, potentially creating harmful pleiotropic effects. The alleles that declined most severely (*Orcu-U1*\*07:01 and *Orcu-U2*\*05:01) had been maintained at high frequencies across all populations, possibly because they provided protection against other pathogens before myxoma virus introduction. Their rapid displacement may increase susceptibility to other infectious diseases or autoimmune conditions, illustrating how adaptation to one pathogen can inadvertently create vulnerabilities to others.

A key outstanding question is the molecular basis of the selective advantage conferred by the MHC alleles under selection. One possibility is that they present specific viral peptides that confer strong protective immunity—a mechanism well established in resistance to RNA viruses^51,52^. Whether this applies to large DNA viruses such as MYXV, which encode a broad array of proteins, remains unclear. However, in chickens we have found that an MHC-II molecule associated with resistance to another large DNA virus, MDV, largely present peptides from just four viral proteins^53^. Alternatively, MYXV is known to suppress MHC surface expression, suggesting that selection may favour alleles that evade or resist this immune evasion strategy, as has been suggested for cytomegaloviruses in the great apes^54^. In both chickens and humans, pathogen resistance has also been linked to variation in the breadth of peptides presented by MHC molecules and in their surface expression levels^55^, raising the possibility that similar mechanisms contribute to MYXV resistance in rabbits.

The myxoma virus pandemic in rabbits offers a compelling example of rapid, parallel adaptation driven by standing genetic variation. The repeated nature of the evolutionary response across continents, the targeted selection on classical MHC-I genes, and the speed of adaptation illuminate key mechanisms underpinning evolutionary rescue during emerging disease outbreaks. These findings underscore the critical role of immunogenetic diversity in enabling population persistence and highlight the value of integrating historical specimens with contemporary genomic approaches to track evolution in real time. As infectious disease and environmental change continue to reshape ecosystems, insights from the rabbit–myxoma system provide a framework for predicting and managing adaptive responses in natural populations.

## Acknowledgments

We thank the ACT Parks and Conservation rangers, the Commonwealth Scientific and Industrial Research Organisation (CSIRO) Rabbit Team, P. Taggart, K. Patel, S. Campbell, the British Association for Shooting and Conservation (BASC) and rabbit hunters for providing rabbit samples in this study. S.Halabi provided rabbit interferon gamma. S.Alfonso assisted with sequencing. We thank L. Loog and J. Cheng for their generous assistance with the analyses. We appreciate the valuable discussion and insights from T. Lenz regarding this study.

## Author contributions

Conceptualization: Y.Z., J.M.A., M.C., J.K., F.M.J.; Methodology: Y.Z., C.T., J.J., F.M.J.; Investigation: Y.Z., J.P.D., M.L., T.S.; Visualisation: Y.Z, J.M.A.; Funding acquisition: J.K., F.M.J.; Project administration: F.M.J.; Supervision: J.K., F.M.J.; Writing – original draft: Y.Z., F.M.J.; Writing – review & editing: all authors.

## Data and materials availability

Original sequence data are available in the Sequence Read Archive (SRA), https://www.ncbi.nlm.nih.gov/sra (BioProject:PRJNA1027777). Assembled MHC-I allele sequences are deposited in GenBank (PV166916 - PV166994, PV166995 - PV167019). Code and other analysis-related files in this study are available at https://doi.org/10.5281/zenodo.16858098.

## Materials and Methods

### Sample collection and published data

To understand the genetic changes occurred in MHC in the European rabbits (*Oryctolagus cuniculus*) in response to myxoma virus across France, Britain and Australia, we generated new sequencing data rabbit samples from these three populations, collected before and after the introduction of myxomatosis. Extra domestic rabbits were included in sequencing for the purpose of identifying MHC alleles. Published datasets were included in some of our analyses. Detailed information on each sample is provided in data file S1, and description on different datasets can be found below.

#### Modern domestic individuals, and rabbits in Britain and Australia (2020-2022)

To characterise the MHC genes in rabbits, we collected 66 tissue samples (liver, spleen, and ear) from modern rabbits: two from domestic individuals, 27 from wild rabbits in Britain, and 37 from wild rabbits in Australia. In addition, two samples from rabbit kidney (RK13) cell line (one infected by interferon-gamma) were kindly provided by our collaborators.

All the rabbit samples from Britain were obtained with the assistance of certified hunters from the British Association for Shooting and Conservation (BASC). No animals were killed specifically for this research, but rather for pest control purposes. Tissues were collected in the field and immediately stored in liquid nitrogen. When extended dissection times were required or liquid nitrogen was unavailable, tissue samples were preserved in falcon tubes with 80% ethanol and maintained on dry ice. These samples were later converted into cDNA libraries and subjected to PacBio long-read transcriptome sequencing. In addition, DNA was extracted from 48 samples and sequenced using MiSeq amplicon sequencing.

#### RNA-Seq Datasets (2022)

To test tissue-specific expression patterns across MHC-I genes, analysis was performed using three RNA-Seq datasets. Two published datasets were obtained from the NCBI Sequence Read Archive (SRA) under accession numbers PRJNA274427 and PRJNA78323. These publicly available datasets provided 19 samples from heart, aorta, liver, kidney, lung, brain, skeletal muscle, and embryo, all sequenced using Illumina paired-end technology. Along with these, we collected 4 samples from Somerset, Britain, and generated 6 tissue-specific samples of spleen (n=3), skeletal muscle (n=2), and kidney (n=1). Detailed information can be found in table S2.

#### Museum samples in France, Britain, and Australia collected before MYXV (1865-1956)

To investigate the genetic basis of resistance to myxoma virus, we obtained 89 DNA library samples from collaborators. These libraries were prepared from historical samples provided by museums, collected from three populations before the release of myxoma virus (France = 26, UK= 31 and Australia = 32), which were included in a previous study ^18^. These 89 libraries were captured using our newly designed probes and subsequently subjected to sequencing.

To enhance sequencing coverage and increase the power of genotyping, we also used the published sequencing data from ^18^, with the abovementioned 89 museum samples included, and an extra sample of 6 (France = 1 and UK = 5) which lacked in DNA libraries for new capture in this study. The capture was designed based on the annotations of the Orycun 2.0 rabbit reference genome (*Ensembl* release 2.69) ^30^, covering 32.10 Mb (1.17%) of the rabbit genome.

#### Modern samples in France, Britain, and Australia collected after RHDV (2002-2013)

To investigate the genetic basis of resistance to myxoma virus, we collected 77 DNA samples donated by collaborators which were included in other study ^18^. These modern samples were collected from three populations (France = 26, UK = 25, Australia = 26) and were used for amplicon sequencing in our lab. While the exome-capture data of modern individuals generated in ^18^ were not directly incorporated into downstream analyses in this study, variant calling results from non-MHC exomes derived for both museum specimens and modern samples from ^18^ were integrated into genome-wide selection scans.

#### Modern samples in Britain, and Australia collected after MYXV, but before RHDV (1985-1996)

To test whether the selection strength has changed after the introduction of RHDV, we obtained 70 tissue (blood and liver) samples kindly provided by collaborators, collected from Britain (n=51) and Australia (n=19), within the period of post-MYXV but pre-RHDV (1985-1996). The samples were then converted into genomic libraries and amplicon sequenced.

#### Modern samples collected specifically in Britain (2012-2022)

To increase sample size for identifying MHC-I alleles, we obtained an additional 14 liver samples from our collaborator. Combined with the 4 samples collected for RNA sequencing, these 18 rabbit samples from Britain were subjected to amplicon sequencing.

For samples sequenced for this study, nomenclatures were attributed based on the type of sample and location. Each sample is classified by a code of four strings of abbreviations connected by an underscore. These four elements include the states of the sample type (H for historical, M for modern, M2 for post-MYXV, pre-RHDV), the country of origin (AU for Australia, FR for France, and UK for Britain), the state/department or county of that country, and the detailed sample localities, followed by the sample number.

### RNA isolation, cDNA library preparation, probe design and captures for PacBio transcriptome sequencing

#### RNA isolation

For RNA extraction, approximately 30mg of frozen rabbit tissue was homogenised under liquid nitrogen using a Cryo-cup grinder (Biospec Products), quickly transferred to a 1.5ml screw top tube containing 500µl of Trizol reagent (Ambion) and approximately ten 1mm diameter zircon beads and immediately homogenised by shaking for 2 minutes at 30Hz in a Tissue Lyser II homogeniser (Qiagen). Homogenised samples were immediately snap frozen in liquid nitrogen and stored at -70°C until processing. RNA was extracted using a standard procedure: samples were thawed at room temperature, incubated for 5 minutes and centrifuged at 17,000g for 10 minutes at 4°C to pellet debris. 450µl of the supernatant was transferred to a fresh 1.5ml tube. 90µlof chloroform was added and the tubes were shaken vigorously for 15 seconds, after a 3-minute incubation at room temperature the samples were centrifuged at 12,000g for 10 minutes at 4°C. 150µl of the upper aqueous phase was removed to a fresh tube and 425µl of propan-2-ol was added, samples were incubated at room temperature for 10 minutes and centrifuged at 12,000g for 10 minutes at 4°C. The supernatant was removed and 250µl of ice cold 70% ethanol was added, and samples were centrifuged at 12,000g for 3 minutes at 4°C. The ethanol was removed by pipetting, and the RNA pellets were resuspended by adding 15µl of nuclease free water (Ambion) and incubating at 42°C for 5 minutes. 1µl of the RNA samples was diluted 30-fold and then quantified using Qubit RNA kit (Thermofisher Scientific) stored in 2µl aliquots at -70°C. RNA integrity was assessed by RNA nano Bioanalyzer chip (Agilent) after diluting the samples to a concentration of approximately 100ng/µl and only samples with an RNA Integrity Number (RIN) of >7 were used for cDNA synthesis.

#### cDNA library preparation

cDNA was synthesised using whole RNA samples, and the Single Cell cDNA Synthesis kit (New England Biolabs). Each sample was labelled with bar-coded polydT primers during the reverse transcription step (synthesised by IDT – PAGE purified). RNA samples were normalised to 100ng/µl by diluting in nuclease free water and 150ng (1.5µl) was added to 1.2µl nuclease free water (Ambion), 0.8µl of 25mM of each dNTP (Thermofisher) and 1µl of 12uM barcoded reverse transcriptase primer (data file S8). The contents were mixed, briefly centrifuged and incubated at 70°C for 5 minutes before snap chilling on ice. 2.5µl of NEBNext Single Cell RT buffer, 1.5µl nuclease free water and 1µl of NEBNext Single Cell RT enzyme mix was added. After mixing and centrifuging briefly samples were incubated in a thermocycler at 42°C with the hot lid set to 52°C for 75 minutes. Samples were placed on ice, 0.5µl of Iso-Seq Express Template Switching Oligo (PacBio) was added, samples were mixed, centrifuged briefly and incubated for a further 15 minutes at 42°C.

15µl of EB (10mM Tris, pH8) was added to each sample, 25µl of ProNex beads (Promega) were added and mixed thoroughly by pipetting. After an incubation of 5 minutes samples were placed on a magnetic stand, when the supernatant was clear it was carefully removed, and samples were washed twice using 200µl of freshly prepared 80% ethanol. After removing the second ethanol wash, tubes were centrifuge briefly at 1000g, returned to the magnetic stand and the last ethanol trace was removed using a 10µl pipette. The beads were allowed to dry until they became matt in appearance then they were removed from the magnetic stand and 23µl of EB was added, and the beads were thoroughly but carefully resuspended. Samples were incubated at 37°C for 5 minutes, replaced on the magnetic stand and 22.8µl of the cleared supernatant containing the cDNA removed to a fresh tube.

The cDNA was amplified by adding to the purified cDNA 25µl of NEBNext Single Cell cDNA PCR Master Mix, 1µl of NEBNext Single Cell cDNA PCR primer, 1µl of Iso-Seq Express cDNA PCR primer and 0.25µl NEBNext Cell Lysis buffer using the following PCR program: 1 cycle of 98°C for 45s then 14 cycles of 98°C for 10s, 62°C for 15s, 72°C for 3 minutes followed by 1 cycle of 72°C for 5 minutes. Amplified cDNA was purified as before using 47.5µl of ProNex beads and quantified using Qubit HS DNA assay (Thermofisher). The quality of library was assessed using an Agilent DNA HS Bioanalyzer kit (Agilent).

#### Probe design, captures and sequencing

*xGen™ Hyb Panel Design Tool* (https://eu.idtdna.com/pages/tools/xgen-hyb-panel-design-tool) was used to design 120mer overlapping biotinylated hybridisation probes (IDT Lockdown Probes), targeting the annotated MHC-I genes in rabbit OryCun2.0 assembly.

Libraries were multiplexed into pools of 24 or 48 using equimolar amounts to give a total quantity of DNA greater than 500ng. For captures an xGen Hybridisation and wash capture kit was used (Integrated DNA Technologies). 500ng of pooled libraries were combined with 7.5µl of Cot DNA and then purified using a 1.8x volume of ProNex beads. A hybridisation mix that comprised 9.5µl of 2x hybridisation buffer, 3µl of Hybridisation buffer enhancer, 1µl of 1m*M* xGen asym TSO blocking oligo, 1µl of 1m*M* xGen RT-primer-barcode block oligo and 4.5µl of custom designed 1X xGen Lockdown probes (Integrated DNA Technologies) was used to resuspend the ProNex beads containing the pooled libraries and Cot DNA. The beads were incubated for 5 minutes at room temperature then placed on a magnet and 17µl of the cleared supernatant was removed to a new 0.2ml PCR tube and incubated for 30s at 95°C then 4 hours at 65°C. 50µl of Dynabeads M-270 Streptavidin were washed twice in 100µl Bead wash buffer and resuspended in 17µl of Bead resuspension mix comprising 8.5µl of xGen 2X Hybridisation buffer, 2.7µl of xGen Hybridisation buffer enhancer and 5.8µl of Nuclease free water. Hybridised cDNA was added to the capture beads and mixed thoroughly and incubated at 65°C for 45 minutes with gentle mixing every 10 minutes. Beads were washed by adding 100µl of wash buffer pre-heated to 65°C, mixing thoroughly by pipetting and collecting beads using a magnetic rack. Cleared supernatant was removed and 150µl of stringent wash buffer pre-heated to 65°C was used to resuspend the beads and the tube was placed immediately at 65°C and incubated for 5 minutes. After collecting the beads and removing the supernatant a second wash with 65°C stringent wash buffer was performed. A further three washes were performed with 150µl of room temperature wash buffer I, II and III respectively and after the final wash the beads were resuspended in 46µl of EB buffer. Amplification of the captured cDNA was performed by adding 50µl of NEBNext High-Fidelity 2X PCR mix, 2µl of NEBNext Single Cell cDNA PCR primer, 2µl of Iso-Seq Express cDNA PCR primer and 0.5µl of NEBNext Cell lysis buffer. The following PCR program was used: 98°C for 45s; 12 cycles of 98°C for 10s, 62°C for 15s, 72°C for 3 minutes; 72°C for 5 minutes. The amplified captured cDNA was cleaned up using 100µl of ProNex beads and eluted in 50µl of EB. The capture was quantified using the Qubit HS DNA assay kit and quality was assessed using an Agilent Bioanalyzer High Sensitivity assay.

A SMRTbell Express Template Prep Kit 2.0 was used for SMRTbell library construction. DNA damage was repaired by adding 7µl of DNA prep buffer, 0.6µl of NAD, 2µl of DNA damage repair mix v2 and 3.2µl of nuclease free water to 44.2µl of amplified cDNA, mixing well and incubating for 30 minutes at 37°C. End repair and A-tailing was carried out by adding on ice 3µl of End Prep mix mixing and incubating at 20°C for 30 minutes then 65°C for 30 minutes. Overhand adapter ligation was performed by adding 3µl of Overhand Adapter v3, 30µl of ligation mix, 1µl of ligation enhancer and 1µl of ligation additive to the previous reaction, mixing well and incubating for 60 minutes at 20°C. Libraries were cleaned using 95µl of ProNex beads using the standard procedure and eluted in 12µl of EB. 1µl of the sample was used to quantify the library using a Qubit DNA HS kit and 0.5µl was diluted 10-fold and the quality and average fragment size analysed on an Agilent Bioanalyser High Sensitivity DNA chip. Sequencing services were provided by Earlham Institute, Norwich, United Kingdom.

The Barcoded Poly-dT primers used for library synthesis are listed in data file S8.

### Generation of RNA sequencing data

#### RNA extraction

Rabbit spleen (n=3), skeletal muscle (n=2) and kidney (n=2) tissues were collected and snap frozen in liquid nitrogen (table S2). Approximately 50mg of tissue was homogenised under liquid nitrogen using a pestle and mortar and transferred immediately into 0.5ml of Tri-reagent (Sigma #) with approximately 10 x 1mm diameter Zirconium beads (Biospec #11079110zx). The tissue was further homogenised by immediately processing for 2 minutes in a TissueLyser II homogeniser (Qiagen) at 30Hz. Samples were centrifuged for 5 minutes at 12000g at 4°C to pellet the debris and 0.4ml of the supernatant was removed to a fresh tube. 80µl of Chloroform was added, the samples were shaken for 15 seconds and centrifuged for 10 minutes at 12000g at 4°C. 150µl of the upper aqueous was removed to a fresh tube and the RNA was precipitated by adding 375µl of Isopropanol, mixing well and incubating at room temperature for 10 minutes. The samples were centrifuged for 10 minutes at 12000g at 4°C and the supernatant removed and discarded. Samples were washed twice with 70% ethanol, and the pellets were resuspended in 15µl of nuclease free water. The integrity of the RNA was assessed using a Bioanalyser RNA high sensitivity chip, those samples exceeding an RNA integrity number of 7 were deemed suitable for library preparation.

#### Library preparation and sequencing

RNAseq libraries were constructed using the NEBNext ® Ultra ™ II Directional RNA Library Prep Kit for Illumina ® with sample purification beads (New England Biolabs # E7765S) according to the manufacturers’ recommendations. PolyA enrichment of the RNA samples was carried out using the NEBNext Poly(A) mRNA Magnetic Isolation Module (New England Biolabs#E7490S). Briefly, 1µg of total RNA was used for each library. After polyA enrichment the RNA was fragmented for 15minutes at 94°C. After end repair and A-tailing adaptor ligation was carried out, the NEBNext adaptor was diluted 5-fold in Adaptor Dilution Buffer prior to ligation. Dual indices were added to the libraries using 8 cycles of PCR and NEBNext ® Multiplex Oligos for Illumina ® (New England Biolabs # E7600S). The libraries were quantified using Qubit High Sensitivity dsDNA assay kit (ThermoFisher Scientific #Q32851). The size distribution and the presence or absence of adaptor dimer was determined using the TapeStation 4200 instrument (Agilent # Q32851) with D1000 Screentapes (Agilent #5067-5582). All libraries contained less than 1% adaptor dimer contamination so passed quality control criteria. The relative quantity of the libraries was ascertained using qPCR: 3 separate 1:1000 dilutions of each library were made using 0.05% Tween20, 10mM Tris-HCl pH8.0. The concentration of the cluster forming molecules in each dilution was determined using 2µl of the diluted library in a qPCR reaction containing 5µl Sensifast Sybr Hi-Rox 2x master mix (Bioline # BIO92005),1µl of library amplification primer mix and 2µl nuclease free water. The qPCR was carried out on a QuantStudio™ 5 Real-Time PCR System, 384-well plate instrument (ThermoFisher # A28570). The libraries were pooled together in equimolar quantities before diluting to 10nM and submitting for paired-end 2×150bp sequencing at the Cancer Research UK Cambridge Institute genomics facility on an Illumina NovaSeq instrument.

### DNA preparation and sourcing of amplicon sequencing

For pre-RHDV samples collected from Norwich, genomic DNA was prepared from rabbit blood samples using a DNeasy Blood and Tissue Kit (Qiagen, 69504) according to the manufacturer’s recommendations. 100µl of blood was used per extraction. Other DNA samples were kind donations from various collaborators (data file S1).

Primers were designed to amplify three overlapping amplicons of size 589, 593 and 633 base pairs covering a total of 1102 bp of genomic sequence of *Orcu-U1* and 1099 bp of *Orcu-U2* (fig. S15a and table S3). Primers were chosen to target sequences conserved between all modern *Orcu-U1* and *Orcu-U2* alleles that had been identified using PacBio transcript sequencing. Where single SNPs existed for some alleles, degenerate base pairs were incorporated into the primer sequences. Part of exon 1, exons 2 and 3 and introns 1 and 2 of MHC-I *Orcu-U1* and *Orcu-U2* were amplified (fig. S15a). 5’ overhang adapter sequence that enabled the later addition of Illumina indices were included. Three PCR reactions were carried out on each sample using genomic DNA templates isolated from rabbits. Each reaction contained 10µl Q5 2 x PCR master mix (NEB, M0492L), 8.5µl nuclease free water (Thermofisher Scientific, AM9930), 0.5µl of 10µM of each primer and 1µl of template DNA. The following PCR program was used: 98°C for 30s; 29 cycles of 98°C for 10s, 68°C for 20s, 72°C for 30s; 72°C for 120s.

The amplicons were examined and quantified using 1% agarose gel electrophoresis and then pooled using 10µl of each. 25µl of pooled PCR was cleaned using 20µl of KAPA Pure beads (Roche, KK8002) according to the manufacturer’s recommendations. The PCR products were eluted from the beads with 50µl of EB buffer (10mM Tris-HCl pH8.0) and 30µl was removed to a fresh PCR plate. Index PCRs were set up using 10µl of Q5 2x PCR master mix, 4µl of nuclease free water, 2µl of pooled cleaned PCR product and 2µl of each forward and reverse Illumina XT indices. The PCR program was as follows: 95°C for 180s; 8 cycles of 95°C for 30s, 55°C for 30s, 72°C for 30s; 72°C for 300s.

A total of 209 indexed PCR products were analysed using 1% agarose gel electrophoresis and then pooled together using an equal volume of each product. 50µl of the pool was cleaned using 56µl of KAPA Pure beads and eluted from the beads with 27.5µl of EB. The resulting pool was quantified using Qubit High Sensitivity DNA assay (Thermofisher, Q32854) and diluted to a final concentration of 10nM. The pool was sequenced using MiSeq 600-cycles v3 kit 2×300bp paired end reads to yield 15Gb of data.

### Capture and sequencing of MHC-I sequences from historical libraries

Probe design for target enrichment was performed using KAPA HyperCap Probes (Roche Sequencing Solutions). The design targeted regions from both the OryCun2.0 (Ensembl release 2.69) and OryCun3.0 rabbit reference genomes (*31*), including specific MHC alleles from five genes (*Orcu-U1, Orcu-U2, Orcu-U3, Orcu-U4* and *Orcu-U8*) that obtained from PacBio long-reads transcriptome sequencing, and *MC1R* gene and its flanking regions. The exact coordinates of targets are available as a BED file in the data file S9.

KAPA HyperExplore MAX 3Mb T3 probes were prepared for hybridisation by resuspension and aliquoting, with the aliquots stored at -20°C. Historical libraries were organised into 14 pools as detailed in data file S1, aiming to enhance library representation based on previous sequencing data. Each pool contained 6-7 libraries, amounting to 0.5-1.5µg of DNA. For pools exceeding 45µl, volume reduction was achieved via vacuum concentration. Pools were normalised to 45µl, to which KAPA Hybrider Reagent was added, followed by incubation with KAPA HyperPure beads. Post-wash, the DNA was resuspended and subjected to a thorough hybridisation process using a carefully calibrated thermocycler program. Post-hybridisation, DNA capture was executed with specific washes and finally eluted in PCR-grade water.

PCR amplification of the enriched DNA employed KAPA HiFi readymix and oligos, with the program consisting of an initial denaturation, followed by cycles for annealing and extension, concluding with a final extension. Clean-up involved KAPA HyperPure beads, with the capture’s quantification and quality assessment performed using Qubit and Agilent assays, respectively. The resulting captures underwent sequencing on the NovaSeq4000 platform at NovoGene, Cambridge, enabling a comprehensive genomic analysis.

### Read mapping, variant calling and MHC-I typing from long-reads transcripts

Circular Consensus Sequence analyses were conducted on the raw data using the *pbccs* tool to obtain all unique representative circular consensus sequences with the default options: --min-passes 3 –min-rq 0.99. This setting requires more than three full-length subreads to generate a CCS for a zero-mode waveguide (ZMW) with predicted accuracy exceeding 0.99. During this step, only reads containing both 5’ and 3’ end primers, as well as the poly-A tail, were retained. The *pbindex* tool was then used to index the resulting bam files. Subsequently, the reads were demultiplexed using the *lima* tool with parameters --isoseq --peek 50000 --guess 75 --guess-min-count 70, based on sample-specific barcodes. The *refine* program from *isoseq 3* was employed to remove poly-A tails and concatemers. All the programs used above for PacBio sequencing data processing were from PacBio Secondary Analysis Tools on Bioconda (https://github.com/PacificBiosciences/pbbioconda). Reads from each individual were then mapped to the rabbit reference genome OryCun 2.0 using *MiniMap2* ^56^ with parameters -ax splice -uf -G 10000. Variant calling and phasing were performed on reads with a quality score greater than 30 using *DeepVariant* ^57^ and *WhatsHap* ^58^.

Genes with >10X sequencing depth for more than half of the individuals were included for our study. The chromosomal locations of these genes, as well as their allele sequences, were meticulously validated using the *Integrated Genomics Viewer (IGV)* ^59^ according to the following criteria: 1)For the quintet of MHC genes incorporated within the capture design, a baseline read count of 10 at each respective locus was anticipated; for the additional paralogs, a minimum count of 5 was acceptable. 2)Verification of a unique allele sequence mandated the presence of at least two reads. 3)The second most prevalent allele was required to exhibit a read count greater than 20% of the one of dominant allele, thereby ensuring allelic heterozygosity was accurately represented.

### Bioinformatic analysis on RNA sequencing data

Following this filtering step, the remaining high-quality reads were aligned to the OryCun2.0 reference genome using *STAR* aligner ^60^ with default parameters. The aligned BAM files were subsequently processed to generate count files. These count files were produced by mapping the aligned reads in the BAM files to annotated gene regions, resulting in counts files in text format. Each count file contained the number of reads that mapped to each gene, necessary for calculating expression levels. The genome annotation file (GTF) used in this step was manually curated to include all the manually characterised MHC Class I genes. The analyses were then performed using *edgeR* package ^61^ in R.

### MHC-I allele typing from amplicon sequencing data

Sequencing reads from 209 rabbit samples were demultiplexed using their unique barcodes. Then, the reads for each sample were segregated based on primer sequences using *Cutadapt* ^62^, requiring these sequences to be anchored at the 5’ end, along with parameters: --max-n 0.1 --minimum-length 100 --trimmed-only --match-read-wildcards --no-indels --error-rate 0. Subsequently, *BBDuk* (http://jgi.doe.gov/data-and-tools/bb-tools/) was then utilised to qualify the reads associated with each primer, excising bases below a quality threshold of 15 from the 3’ end and discarding reads whose average quality below 15.

The downstream pipeline was illustrated in fig. S15b. According to the per base sequence quality report by *FASTQC* ^63^, a notable decline in quality scores at approximately 150 base pairs was observed across the majority of individuals, which was consistent with the patterns in chicken sequences (Clive Tregaskes, Rebecca Martin and Jim Kaufman, unpublished). Consequently, the first 130 bp from R1 and R2 sequences were concatenated to form an index and reads shorter than 130 base pairs were excluded from further analysis.

These indices were then clustered by sequence identity, and major allele groups were defined as those supported by a minimum percentage of the total reads (e.g., 1%). To mitigate potential discrepancies in allele group assignment, an extended index was generated from longer reads to resolve closely related alleles. The length for these reads was determined by the 70th percentile of read lengths within the primary cluster. This secondary clustering created sub-groups which were retained if they represented at least 10% of the reads and were supported by a minimum of 5 reads.

For each defined allele group, the top 100 longest reads were aligned using MAFFT ^64^. A custom consensus sequence was then generated, requiring a base to be present in at least 75% of the reads at a given position to be called; otherwise, it was marked as an ‘X’. Gaps present in over 75% of reads were retained in the consensus. These consensus sequences were clipped to exon boundaries, the R2-derived sequence was reverse-complemented, and the parts were concatenated.

The resulting consensus sequences from all three amplicons were aligned together using MAFFT ^64^ and deduplicated. Primer sequences were used as anchors to identify overlapping regions between amplicons (amplicon 1 with 3, and amplicon 3 with 2). Pairs of sequences with 100% identity in their overlapping region were merged. This process was repeated to assemble full-length sequences spanning from exon 1 to exon 3.

Assembled full-length sequences were validated by comparing them against a curated database of known alleles derived from previous PacBio sequencing. Novel alleles not found in the database were subsequently checked for chimerism against the successfully validated alleles.

To ensure the robustness of our results, we performed a sensitivity analysis by running the pipeline with three distinct parameter sets. This involved varying the minimum read support threshold for amplicon 2 (0.5% vs. 1.0%) and executing the analysis both with and without an updated allele reference database. A multi-step filtering protocol was then applied to the resulting alleles from each run. First, to remove potential assembly artifacts, any allele was discarded if its read support was less than 50% of the most abundant allele assembled from the same set of amplicons. Second, to establish a final confidence threshold for genotyping, an allele was removed if its support was less than 25% of the most abundant validated allele within that sample. Finally, to maximise data recovery, partial alleles constructed from a successful amplicon 1-and-3 assembly were retained in cases where a full-length assembly failed.

### Genomic map of MHC region

The MHC region annotations for chr12: 20,000,000-24,000,000 were obtained from *Ensembl* rabbit OryCun 2.0 reference genome assembly. The coordinates of the nine MHC-I paralogs (*Orcu-U1* to *Orcu-U9)* were curated using PacBio sequencing reads (as described above). MHC-II paralogs were also curated from the same dataset but are not discussed in this research (Marina Lirintzi *et. al*; unpublished). The plot includes MHC Class III genes and several conserved genes in MHC region across species, encoding molecules involved in peptide-binding and peptide editing ^19^, based on *Ensembl* annotations. Genes included in this analysis is recorded in data file S10.

### MUMmer analyses

The nucleotide sequence spanning 0.14 Mb, specifically from positions chr12: 20,180,000-20,320,000 within the MHC-I region of the OryCun 2.0 reference genome assembly was analysed to ascertain the presence of genomic duplications. This examination was conducted using the software *MUMmer* ^65^. The analysis was parameterised with -maxmatch -b -c -l 50 to identify repeat sequences of 50 base pairs or more, visualised using a dot plot. The same analysis using identical parameter set was conducted for the region containing the second cluster of MHC-I genes (chr12: 22,300,000-22,600,000).

### Nucleotide diversity and non-synonymous genetic diversity

Nucleotide diversity and associated non-synonymous genetic variation within the signal peptide, α1, α2, and α3 domains of each gene were quantified. Correlated statistics of synonymous and non-synonymous genetic diversity for each domain were subsequently produced. These were then complemented by assessments of haplotype diversity, Tajima’s D, Watterson’s Theta, and the nucleotide diversity encompassing the entire gene regions. All the aforementioned analyses were performed using the *ape* ^66^ and *pegas* ^67^ package in R, with the 64 individuals categorised into UK and Australian populations. 95% bootstrap confidence intervals (CIs) were estimated with 1,000 replacements within each population.

To generate a genome-wide baseline for comparison, mean genome-wide nucleotide diversity was calculated for the modern UK and Australian populations using whole-exome sequencing data derived from identical samples in a previous study ^18^. For each population, per-site diversity was first calculated for all variant sites using VCFtools with the *--site-pi* flag. To correctly account for invariant sites, the sum of these per-site π values was then divided by the total number of base pairs in the exome capture target design, yielding a single average value for each population.

### Phylogenetic analyses

Maximum likelihood phylogeny of MHC and MHC-I-like genes of rabbits and humans was reconstructed based on alignment of 48 amino acid sequences using *IQ-TREE* ^68^. GTR20 was used to model evolutionary rate differences among sites. Branch support is indicated as percentage of trees out of 1,000 bootstrap replicates. The genes with names starting as *Orcu* are characterised based on the PacBio transcriptome sequencing data, while the ones starting by R are the rabbit paralogs annotated by *Ensembl*. HLA-like paralogs included are adapted from ^69^. Protein sequences alignment included for this phylogeny can be found in data file S4.

Domain-specific phylogenetic analyses were conducted on all alleles from the nine candidate MHC-I genes, separating the SP, α1, α2, and α3 domains. The nucleotide sequences of each domain were aligned using *MAFFT* ^64^, and then phylogeny was constructed using JC69 model with 1,000 bootstrap replications using *IQ-TREE* ^68^. The clades of the trees were coloured according to different genes, using *ggtree* v3.10.1 ^70^.

The phylogeny including the 29 alleles from gene *Orcu-U1* was performed with JTT amino acid substitution model using *IQ-TREE* ^68^.

### Nomenclature of MHC Class I alleles

To name the identified alleles, we followed an universal nomenclature criteria for non-human MHC genes ^71^. The MHC symbol is followed by a four-letter abbreviation of the species scientific name, thus the MHC-I genes of *Oryctolagus cuniculus* would was named accordingly as *Orcu.* Then a hyphen divides the four-letter designation from the gene name, designated by capital letters and Arabic numerals. The gene names were given based on the level of polymorphism, the potential of being a classical MHC Class I genes, and the order of their initial discovery.

In this study, a total of 104 alleles encompassing nine MHC-I genes were identified. 79 alleles were identified through PacBio sequencing, while the remaining 25 alleles, specific to *Orcu*-*U1* and *Orcu*-*U2* genes, were delineated via MiSeq amplicon sequencing. The MiSeq sequencing facilitated the assembly of sequences spanning 561 base pairs, which included portions of exon 1 and 3, as well as the full sequence of exon 2. Nonetheless, this sequence collection was challenged by a varying number of unidentified nucleotide bases, predominantly localised at the juncture of exon 1 and exon 2, with the maximum gap being 33 base pairs in length.

Employing the 528 base pair sequences of exon 2 and 3, which translate to 176 amino acids of peptide binding domains, a distance matrix was generated for every allele pair within each gene (fig. S4). To classify allele groups, a threshold was set at eight amino acid discrepancies. Alleles that did not aggregate into existing groups were designated into new allele groups. In situations involving alleles with ambiguous residues, comparisons were conducted to prevent adding any mismatch assignment from these sites. Therefore, the first two series of digits in the nomenclature denote the extent of amino acid variation within this region. The third set of digits signifies synonymous nucleotide substitutions occurring within the same segment, while the final set of digits corresponds to variances beyond the regions spanned by the two exons, taking any diversity from other regions into consideration. The short nomenclatures have been created by omitting digit sets from the fourth to the second position, only if the resulting names remain unique.

### MHC-I allele typing from historical samples

The MHC-I region, characterised by an extraordinarily high level of polymorphism and the fragmented nature of historical DNA, presents significant challenges in accurately calling variants from Illumina short reads derived from historical samples. To address this, we adapted the published tool *OptiType* ^33^, initially designed for non-novel HLA allele genotyping from NGS data through integer linear programming. The utility of *OptiType* in our context optimise the capacity to align multiple allele sequences across multiple loci, subsequently optimising the genotype configuration that accounts for the maximum number of mapped reads.

For input, *OptiType* requires a collection of known MHC-I alleles; our dataset comprised 97 allele sequences spanning *Orcu-U1* to *Orcu-U8*, compiled from our preceding sequencing efforts. *Orcu-U9* was excluded from this and downstream analyses due to length variations in the α1 and α2 domains relative to other paralogs. To mitigate potential reference sequence length biases during allele calling, we standardised all reference sequences to the exon 1-3 region. This truncation corresponds with the target region of the MiSeq sequencing, encompassing a 561 bp stretch.

To rigorously evaluate the precision of our methodology, we conducted a simulation analysis, utilising all 79 unique haplotype combinations identified in *Orcu-U1* and *Orcu-U2* from modern and pre-RHDV specimens. We employed *Gargammel* ^72^ to generate Illumina short-read simulations based on these haplotype patterns, assigning alleles from the remaining seven paralogs randomly to preclude any biases.

Our simulation parameters were calibrated to mimic real historical data: fragment lengths were set to 40 bp, and read lengths to 100 bp, inclusive of post-mortem damage patterns reflective of actual historical samples. Various parameter sets were tested to determine their optimisation for downstream analysis—coverage levels spanned from 0.5X to 50X, heterozygosity bias ranged from 0 to 0.1 in step of 0.005, and the mismatch tolerance for aligner was adjusted among match=90%, match=95%, match=97%, and match=100%. Acknowledging uncertainties in calling alleles from the seven additional paralogs, we generated simulated reads for each haplotype pair across 100 bootstrap replicates, and the average accuracy was calculated to reflect the requirement of dataset, as well as the optimal parameter sets of *OptiType* in genotyping *Orcu-U1* and *Orcu-U2* alleles.

The sequencing data from our new captures of museum samples targeting MHC, as well as the published old capture data ^18^ were combined to enhance the power and optimise the level of coverage. Of the 95 individuals, only 79 met the required coverage threshold of >3X and were included in the downstream analyses, accounting for the accuracy requirements of *OptiType* genotyping (data file S1). For these individuals, adapters and poly-A tails were trimmed, followed by allele typing using *OptiType*.

### Changes in allele frequency

We investigated the allelic frequency variations before and after the myxoma virus introduction by analysing a historical cohort and a modern one, composing up to 172 individuals from three rabbit populations. The historical cohort comprised 95 individuals, with a mean sequencing depth of 16X. A subset of 79 samples achieved adequate coverage (>3X), classified by population France = 30, UK = 32, Australia = 17 (data file S1). The coverage metrics were calculated based on *MC1R* gene, which was included in our capture design (data file S1). 77 modern samples (France = 26, UK = 25, Australia = 26), coinciding with the geographic collection points of the historical specimens, were included in this analysis with their alleles assembled from previous MiSeq amplicon sequencing.

Allelic frequencies for *Orcu-U1* and *Orcu-U2* loci were computed across the three populations at two temporal points. Historical allele calling employed *OptiType*, with quality delineated by mapping depth across a 561 base-pair region and site-specific depth occurrence. Alleles with an average mapping depth exceeding 3 and over 400 sites of non-zero depth were designated ‘high-quality’. Those not meeting these criteria were classified as ‘low-quality’. We plotted the frequency distributions of alleles from both *Orcu-U1* and *Orcu-U2* genes, contrasting historical and modern datasets for each population, with the low-quality calls of alleles from historical individuals integrated as stacked grey bars (fig. S10).

### Selection scan comparing modern and historical samples

The *Ohana* software ^34^ was utilised to infer the covariance structure in the MHC-I allele frequencies across three populations. In analysing the genome-wide variants, we relied on genotype data derived from whole exome sequencing, which has been produced by previous study ^18^ for identical samples to our study. The variants for whole exome region were collected from a filtered VCF ^18^, and only variants with a minimum of 10 genotypes and a minor allele frequency of 0.05 across all individuals were kept, yielding a final count of 756,482 variants. To ascertain the variant details for *Orcu-U1* and *Orcu-U2* within the within the MiSeq amplicon sequencing region, we performed variant calling using the coding sequences from both modern and historical individual samples. We removed the 58 variants existing in the original VCF and added 123 newly called variants from the coding regions of *Orcu-U1* and *Orcu-U2*. It is noteworthy that any historical allele variants with zero depth in the *OptiType* calls were annotated as missing genotypes in the VCF. Only those variants demonstrating a minor allele frequency (MAF) of 0.05, which was consistent to the threshold applied to other genomic regions, were included in the newly constructed VCF. Subsequently, our analysis was confined to individuals present in both datasets, resulting in 150 samples (Historical = 75: France = 29, UK = 29, Australia = 17; Modern = 75: France = 25, UK = 24, Australia = 26), with 756,547 variants being incorporated into the final Ohana analyses.

In our selection scan analysis, we employed a time-aware modelling approach which accommodates temporally labelled data, enabling consideration of changes in allele frequencies that are dependent on specific time frames. We used the dates that were historically documented which marking the initial introduction of the virus into each country as one of the inputs for the model (i.e. 1952 for France, 1953 for Britain, and 1950 for Australia). The resulting likelihood ratios for each genetic variant were then visualised, offering both genome-wide and MHC-I region-specific perspectives. Detailed Ohana models can be found in the supplementary file in ^18^.

To establish a genome-wide 95% significance threshold, we conducted a permutation analysis. This required shuffling the year of sample collection within each country, and repeating the above analyses for 1,000 iterations, during each of which the maximum likelihood ratio statistic was recorded. These permuted datasets provided a robust basis for determining the significance threshold against which the observed likelihood ratios were compared.

### Detection of changes in selection strength

We employed a Bayesian approach to infer the timing and strength of selection acting on MHC haplotypes, adapting the method described in ^42^.

Genotypes of 299 individuals, from modern (n=79) and historical (n=150), and pre-RHDV (n=70) time periods, across three populations (data file S1) were phased using *PHASE* version 2.1 ^37^. Phased haplotype data was processed to identify common haplotypes (represented by ≥5 individuals) to include for the downstream analysis. Subsequently, the output was converted into binary format, where each haplotype was classified as either the target haplotype or “other” for analysis.

The likelihood of observing the data was calculated as the product of probabilities for all observed haplotypes across samples, incorporating both haplotype frequency at each time points and sample age. Haplotype frequency trajectories were modelled using a standard additive selection model (h = 0.5), where frequency changes depend on the selection coefficient and current haplotype frequency.

We implemented a time-dependent selection model with two phases. Selection was assumed to begin with myxoma virus introduction (1950 in Australia, 1953 in UK) at coefficient s₁, then change upon RHDV arrival (1995 in Australia, 1992 in UK) to coefficient s₂. Before myxoma virus introduction, haplotype frequencies remained constant at ancestral levels.

Parameter estimation involved comprehensive parameter sweeps across uniform priors for three key variables: ancestral haplotype frequency (0-1, steps of 0.01) and two selection coefficients s₁ and s₂ (ranging from -1 to 1, with steps of 0.01-0.005). For each parameter combination, we calculated deterministic allele frequency trajectories using the *lsoda* function in R package *deSolve* ^73^. Marginal posterior distributions were obtained by numerically integrating likelihoods over the remaining parameters. Confidence intervals were calculated from likelihood surfaces, and the significance of selection coefficient changes (s₂-s₁) was assessed using standard error propagation methods.

### Heterozygosity and allele divergence

Expected heterozygosity (*He*) was calculated for each population-time-gene combination using the standard formula: *He* = 1 - Σpi², where pi represents the frequency of the i-th allele. *He* values were calculated from observed allele frequencies in the real data, representing the expected frequency of heterozygotes under Hardy-Weinberg equilibrium. Confidence intervals for He estimates were calculated using bootstrap resampling with 1,000 iterations. For each bootstrap replicate, individuals were resampled with replacement within each population-time-gene combination, maintaining the original sample size. *He* was recalculated for each bootstrap replicate, and 95% confidence intervals were estimated from the bootstrap distribution. Significance of differences in He between historical and modern samples was assessed using permutation tests with 10,000 iterations. For each permutation, time period labels (Historical vs Modern) were randomly reassigned to individuals within each population-gene combination, while maintaining the original sample sizes for each time period. *He* values were recalculated for the permuted groups, and the difference between permuted historical and modern *He* values was recorded. The empirical p-value was calculated as the proportion of permuted differences with absolute values greater than or equal to the observed difference. P-values < 0.05 were considered statistically significant.

Expected mean allele divergence was calculated for each population-time-gene combination as the Hardy-Weinberg weighted average of pairwise Grantham distances between all alleles. For each population-time-gene combination, we calculated the expected mean divergence between alleles in heterozygotes using the formula: Expected Mean Divergence = Σ(2 × pi × pj × dij) / Σ(2 × pi × pj), where pi and pj represent the frequencies of alleles i and j, and dij represents the Grantham distance between them. The weights (2 × pi × pj) represent the probability of each heterozygote combination under Hardy-Weinberg equilibrium.

Confidence intervals were calculated using bootstrap resampling with 1,000 iterations, resampling individuals with replacement within each population-time-gene combination. Significance of differences between historical and modern samples was assessed using permutation tests with 1,000 iterations, where time period labels were randomly reassigned to individuals while maintaining original sample sizes.

To test whether alleles with higher frequencies tend to be more divergent from other common alleles, we calculated the average Grantham pairwise divergence for each allele to all common alleles (≥5% frequency) within the same population and gene. For each allele, we computed the mean Grantham distance to all other common alleles with frequency ≥5% in the same population-gene combination, representing the most likely heterozygote partners. The analysis included all alleles with frequency ≥1% to ensure adequate sample size. We then tested for correlations between this average divergence measure and allele frequency using Spearman’s rank correlation coefficient. Statistical significance was assessed at α = 0.05, with results reported as significant (*p<0.05) or non-significant (ns)*.

## References

1. Golas, B. D., Goodell, B. & Webb, C. T. Host adaptation to novel pathogen introduction: Predicting conditions that promote evolutionary rescue. Ecology Letters 24, 2238–2255 (2021).

2. Barrett, R. D. H. & Schluter, D. Adaptation from standing genetic variation. Trends in Ecology & Evolution 23, 38–44 (2008).

3. Radwan, J., Babik, W., Kaufman, J., Lenz, T. L. & Winternitz, J. Advances in the Evolutionary Understanding of MHC Polymorphism. Trends in Genetics 36, 298–311 (2020).

4. Heijmans, C. M. C., Groot, N. G. & Bontrop, R. E. Comparative genetics of the major histocompatibility complex in humans and nonhuman primates. Int J Immunogenet 47, 243–260 (2020).

5. Moore, C. B. et al. Evidence of HIV-1 Adaptation to HLA-Restricted Immune Responses at a Population Level. Science 296, 1439–1443 (2002).

6. Hedrick, P. W. Pathogen resistance and genetic variation at MHC loci. Evolution 56, 1902–1908 (2002).

7. Eizaguirre, C., Lenz, T. L., Kalbe, M. & Milinski, M. Rapid and adaptive evolution of MHC genes under parasite selection in experimental vertebrate populations. Nat Commun 3, 621 (2012).

8. Carrington, M. et al. HLA and HIV-1: heterozygote advantage and B* 35-Cw* 04 disadvantage. Science 283, 1748–1752 (1999).

9. Trachtenberg, E. et al. Advantage of rare HLA supertype in HIV disease progression. Nat Med 9, 928–935 (2003).

10. Kawashima, Y. et al. Adaptation of HIV-1 to human leukocyte antigen class I. Nature 458, 641–645 (2009).

11. Prugnolle, F. et al. Pathogen-Driven Selection and Worldwide HLA Class I Diversity. Current Biology 15, 1022–1027 (2005).

12. Eizaguirre, C. & Lenz, T. L. Major histocompatibility complex polymorphism: dynamics and consequences of parasite-mediated local adaptation in fishes. Journal of fish biology 77, 2023–2047 (2010).

13. Calvignac-Spencer, S. & Lenz, T. L. The One Past Health workshop: connecting ancient DNA and zoonosis research. BioEssays 39, 1–4 (2017).

14. Plascencia, A. G., Jakobsson, M. & Sánchez-Quinto, F. Ancient DNA HLA typing reveals significant shifts in frequency in Europe since the Neolithic. Sci Rep 15, 6161 (2025).

15. Fenner, F. & Ratcliffe, F. N. Myxomatosis. (Cambridge University Press, Cambridge, UK, 1965).

16. Fenner, F. & Fantini, B. Biological Control of Vertebrate Pests: The History of Myxomatosis, an Experiment in Evolution. xii + 339 pp. (CABI Publishing, Wallingford, 1999).

17. Kerr, P. J. Myxomatosis in Australia and Europe: A model for emerging infectious diseases. Antiviral Research 93, 387–415 (2012).

18. Alves, J. M. et al. Parallel adaptation of rabbit populations to myxoma virus. Science 363, 1319–1326 (2019).

19. Kulski, J. K., Suzuki, S. & Shiina, T. Human leukocyte antigen super-locus: nexus of genomic supergenes, SNPs, indels, transcripts, and haplotypes. Hum Genome Var 9, 1–15 (2022).

20. Rodgers, J. R. & Cook, R. G. MHC class Ib molecules bridge innate and acquired immunity. Nat Rev Immunol 5, 459–471 (2005).

21. D’Souza, M. P. et al. Casting a wider net: Immunosurveillance by nonclassical MHC molecules. PLOS Pathogens 15, e1007567 (2019).

22. Dalbey, R. E., Lively, M. O., Bron, S. & Dijl, J. M. V. The chemistry and enzymology of the type I signal peptidases. Protein Science 6, 1129–1138 (1997).

23. Seidel, E. et al. A slowly cleaved viral signal peptide acts as a protein-integral immune evasion domain. Nat Commun 12, 2061 (2021).

24. Ilca, F. T. & Boyle, L. H. The glycosylation status of MHC class I molecules impacts their interactions with TAPBPR. Molecular Immunology 139, 168–176 (2021).

25. Lan, H. et al. Exchange catalysis by tapasin exploits conserved and allele-specific features of MHC-I molecules. Nat Commun 12, 4236 (2021).

26. Park, B. et al. The Truncated Cytoplasmic Tail of HLA-G Serves a Quality-Control Function in Post-ER Compartments. Immunity 15, 213–224 (2001).

27. Lizée, G., Basha, G. & Jefferies, W. A. Tails of wonder: endocytic-sorting motifs key for exogenous antigen presentation. Trends in Immunology 26, 141–149 (2005).

28. Basha, G. et al. MHC Class I Endosomal and Lysosomal Trafficking Coincides with Exogenous Antigen Loading in Dendritic Cells. PLoS One 3, e3247 (2008).

29. Madden, D. R., Gorga, J. C., Strominger, J. L. & Wiley, D. C. The structure of HLA-B27 reveals nonamer self-peptides bound in an extended conformation. Nature 353, 321–325 (1991).

30. Carneiro, M. et al. Rabbit genome analysis reveals a polygenic basis for phenotypic change during domestication. Science 345, 1074–1079 (2014).

31. Hughes, A. L. & Nei, M. Evolution of the major histocompatibility complex: independent origin of nonclassical class I genes in different groups of mammals. Molecular Biology and Evolution 6, 559–579 (1989).

32. Pease, L. R., Schulze, D. H., Pfaffenbach, G. M. & Nathenson, S. G. Spontaneous H-2 mutants provide evidence that a copy mechanism analogous to gene conversion generates polymorphism in the major histocompatibility complex. Proceedings of the National Academy of Sciences 80, 242–246 (1983).

33. Szolek, A. et al. OptiType: precision HLA typing from next-generation sequencing data. Bioinformatics 30, 3310–3316 (2014).

34. Cheng, J. Y., Stern, A. J., Racimo, F. & Nielsen, R. Detecting Selection in Multiple Populations by Modeling Ancestral Admixture Components. Molecular Biology and Evolution 39, msab294 (2022).

35. Veale, E. M. The Rabbit in England. The Agricultural History Review 5, 85–90 (1957).

36. Alves, J. M. et al. A single introduction of wild rabbits triggered the biological invasion of Australia. Proceedings of the National Academy of Sciences 119, e2122734119 (2022).

37. Stephens, M., Smith, N. J. & Donnelly, P. A New Statistical Method for Haplotype Reconstruction from Population Data. The American Journal of Human Genetics 68, 978–989 (2001).

38. Xu, Z. J. & Chen, W. X. Viral haemorrhagic disease in rabbits: A review. Vet Res Commun 13, 205–212 (1989).

39. Abrantes, J., van der Loo, W., Le Pendu, J. & Esteves, P. J. Rabbit haemorrhagic disease (RHD) and rabbit haemorrhagic disease virus (RHDV): a review. Veterinary Research 43, 12 (2012).

40. Morisse, J. P., Le Gall, G. & Boilletot, E. Hepatitis of viral origin in Leporidae: introduction and aetiological hypotheses. Rev Sci Tech 10, 269–310 (1991).

41. Fuller, H. E., Chasey, D., Lucas, M. H. & Gibbens, J. C. Rabbit haemorrhagic disease in the United Kingdom. Vet Rec 133, 611–613 (1993).

42. Loog, L. et al. Inferring Allele Frequency Trajectories from Ancient DNA Indicates That Selection on a Chicken Gene Coincided with Changes in Medieval Husbandry Practices. Molecular Biology and Evolution 34, 1981–1990 (2017).

43. Potts, W. K. & Wakeland, E. K. Evolution of diversity at the major histocompatibility complex. Trends in Ecology & Evolution 5, 181–187 (1990).

44. Wakeland, E. K. et al. Ancestral polymorphisms of MHC class II genes: divergent allele advantage. Immunol Res 9, 115–122 (1990).

45. Krause-Kyora, B. et al. Ancient DNA study reveals HLA susceptibility locus for leprosy in medieval Europeans. Nature Communications 9, (2018).

46. Ross, J. & Sanders, M. F. The development of genetic resistance to myxomatosis in wild rabbits in Britain. Epidemiology & Infection 92, 255–261 (1984).

47. Fenner, F. & Ross, J. Myxomatosis. in The European Rabbit: The history and biology of a successful colonizer (eds Thompson, H. V. & King, C. M.) 0 (Oxford University Press, 1994). doi:10.1093/oso/9780198576112.003.0007.

48. Marshall, I. D. & Fenner, F. Studies in the epidemiology of infectious myxomatosis of rabbits. Epidemiology & Infection 56, 288–302 (1958).

49. Cooke, B. Long-term monitoring of disease impact: rabbit haemorrhagic disease as a biological control case study. Vet Rec 182, 571–572 (2018).

50. Elsworth, P. G., Kovaliski, J. & Cooke, B. D. Rabbit haemorrhagic disease: are Australian rabbits (Oryctolagus cuniculus) evolving resistance to infection with Czech CAPM 351 RHDV? Epidemiology & Infection 140, 1972–1981 (2012).

51. Woolthuis, R. G., van Dorp, C. H., Keşmir, C., de Boer, R. J. & van Boven, M. Long-term adaptation of the influenza A virus by escaping cytotoxic T-cell recognition. Sci Rep 6, 33334 (2016).

52. Arcia, D., Acevedo-Sáenz, L., Rugeles, M. T. & Velilla, P. A. Role of CD8+ T Cells in the Selection of HIV-1 Immune Escape Mutations. Viral Immunol 30, 3–12 (2017).

53. Halabi, S. et al. The dominantly expressed class II molecule from a resistant MHC haplotype presents only a few Marek’s disease virus peptides by using an unprecedented binding motif. PLoS Biol 19, e3001057 (2021).

54. Zimmermann, C. et al. Diverse cytomegalovirus US11 antagonism and MHC-A evasion strategies reveal a tit-for-tat coevolutionary arms race in hominids. Proc. Natl. Acad. Sci. U.S.A. 121, e2315985121 (2024).

55. Chappell, P. et al. Expression levels of MHC class I molecules are inversely correlated with promiscuity of peptide binding. eLife 2015, 1–22 (2015).

56. Li, H. Minimap2: pairwise alignment for nucleotide sequences. Bioinformatics 34, 3094– 3100 (2018).

57. Poplin, R. et al. A universal SNP and small-indel variant caller using deep neural networks. Nat Biotechnol 36, 983–987 (2018).

58. Patterson, M. D. et al. WhatsHap: Weighted Haplotype Assembly for Future-Generation Sequencing Reads. Journal of Computational Biology 22, 498–509 (2015).

59. Thorvaldsdóttir, H., Robinson, J. T. & Mesirov, J. P. Integrative Genomics Viewer (IGV): high-performance genomics data visualization and exploration. Brief Bioinform 14, 178–192 (2013).

60. Dobin, A. et al. STAR: ultrafast universal RNA-seq aligner. Bioinformatics 29, 15–21 (2013).

61. Robinson, M. D., McCarthy, D. J. & Smyth, G. K. edgeR: a Bioconductor package for differential expression analysis of digital gene expression data. Bioinformatics 26, 139 (2009).

62. Martin, M. Cutadapt removes adapter sequences from high-throughput sequencing reads. EMBnet.journal 17, 10–12 (2011).

63. Andrews, S. FastQC: a quality control tool for high throughput sequence data. http://www.bioinformatics.babraham.ac.uk/projects/fastqc (2010).

64. Katoh, K., Misawa, K., Kuma, K. & Miyata, T. MAFFT: a novel method for rapid multiple sequence alignment based on fast Fourier transform. Nucleic Acids Research 30, 3059–3066 (2002).

65. Marçais, G. et al. MUMmer4: A fast and versatile genome alignment system. PLOS Computational Biology 14, e1005944 (2018).

66. Paradis, E., Claude, J. & Strimmer, K. APE: Analyses of Phylogenetics and Evolution in R language. Bioinformatics 20, 289–290 (2004).

67. Paradis, E. pegas: an R package for population genetics with an integrated–modular approach. Bioinformatics 26, 419–420 (2010).

68. Nguyen, L.-T., Schmidt, H. A., von Haeseler, A. & Minh, B. Q. IQ-TREE: A Fast and Effective Stochastic Algorithm for Estimating Maximum-Likelihood Phylogenies. Mol Biol Evol 32, 268–274 (2015).

69. Maruoka, T., Tanabe, H., Chiba, M. & Kasahara, M. Chicken CD1 genes are located in the MHC: CD1 and endothelial protein C receptor genes constitute a distinct subfamily of class-I-like genes that predates the emergence of mammals. Immunogenetics 57, 590–600 (2005).

70. Yu, G., Smith, D. K., Zhu, H., Guan, Y. & Lam, T. T.-Y. ggtree: an r package for visualization and annotation of phylogenetic trees with their covariates and other associated data. Methods in Ecology and Evolution 8, 28–36 (2017).

71. Maccari, G. et al. IPD-MHC: nomenclature requirements for the non-human major histocompatibility complex in the next-generation sequencing era. Immunogenetics 70, 619–623 (2018).

72. Renaud, G., Hanghøj, K., Willerslev, E. & Orlando, L. gargammel: a sequence simulator for ancient DNA. Bioinformatics 33, 577–579 (2017).

73. Soetaert, K., Petzoldt, T. & Setzer, R. W. Solving Differential Equations in R: Package deSolve. J. Stat. Soft. 33, (2010).

