## Supplementary Materials for "MHC-I diversity enables rapid adaptation during viral pandemic in wild rabbit populations"

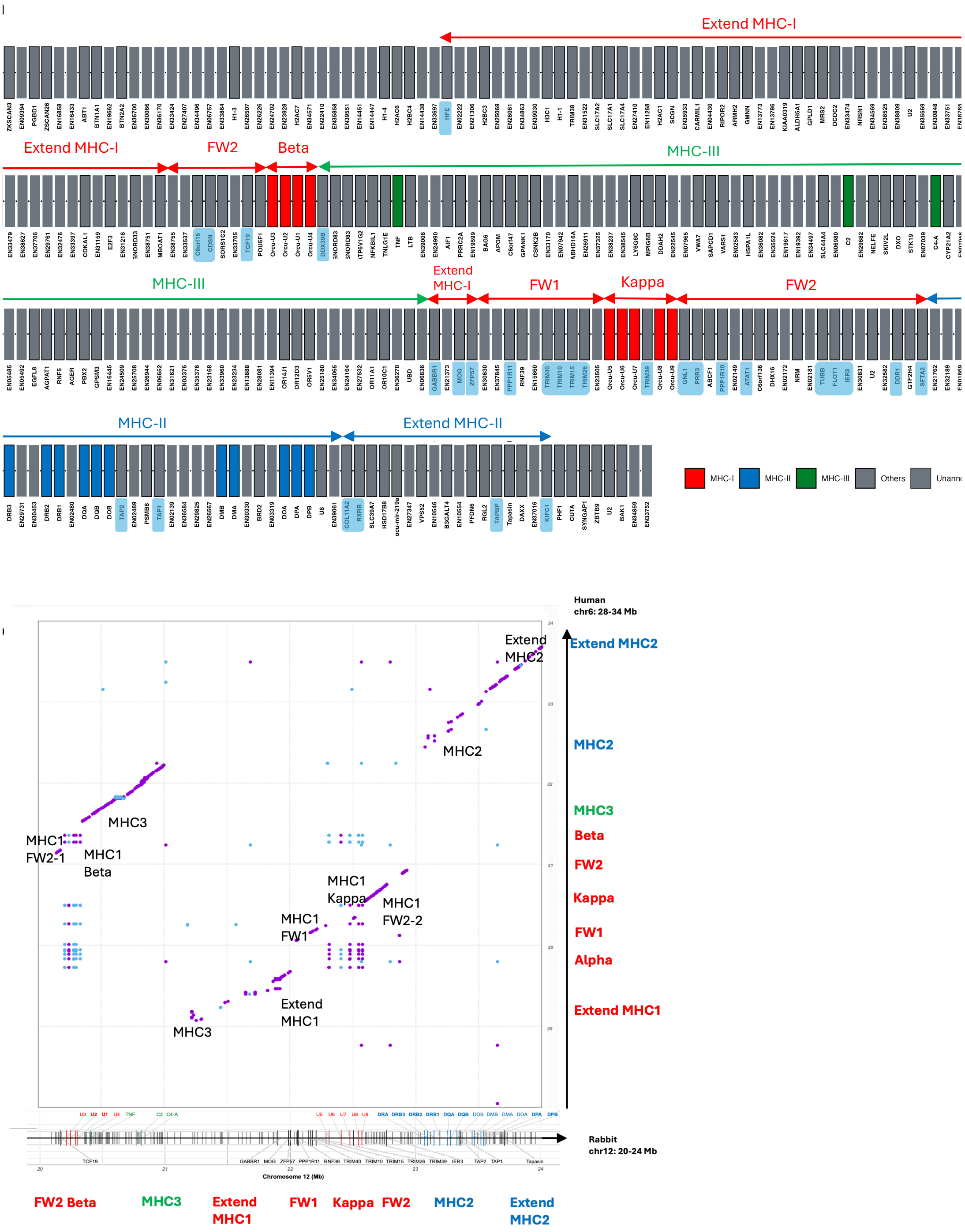


**Fig. S1. chromosomal rearrangement of the MHC region in rabbits. (a)** Physical genomic map of the rabbit MHC region on chromosome 12 (OryCun2.0, 10.5-24 Mb). Gene positions correspond to physical locations on chromosome 12 with continuous composition from left to right side (3 rows). Genes were either annotated within assembly or newly annotated from this study (names starting with '*Orcu*'). Names starting with 'EN' indicate unannotated genes from assembly. Coloured boxes identify MHC-I, MHC-II and MHC-III genes, with shaded gene names representing genes identified also in human MHC region. Rabbit MHC-I region is organised into distinct blocks: Beta, Extend MHC-I, Kappa, accompanied with framework (FW) regions containing non-MHC genes. **(b)** Dot plot comparing rabbit chr12: 20-24Mb (x-axis) with human chr6: 28-34Mb (y-axis), showing conserved syntenic blocks and the chromosomal rearrangement in rabbit MHC region. Dots represent regions >100bp with over 80% similarity, with two different colours indicating different directions of the match.

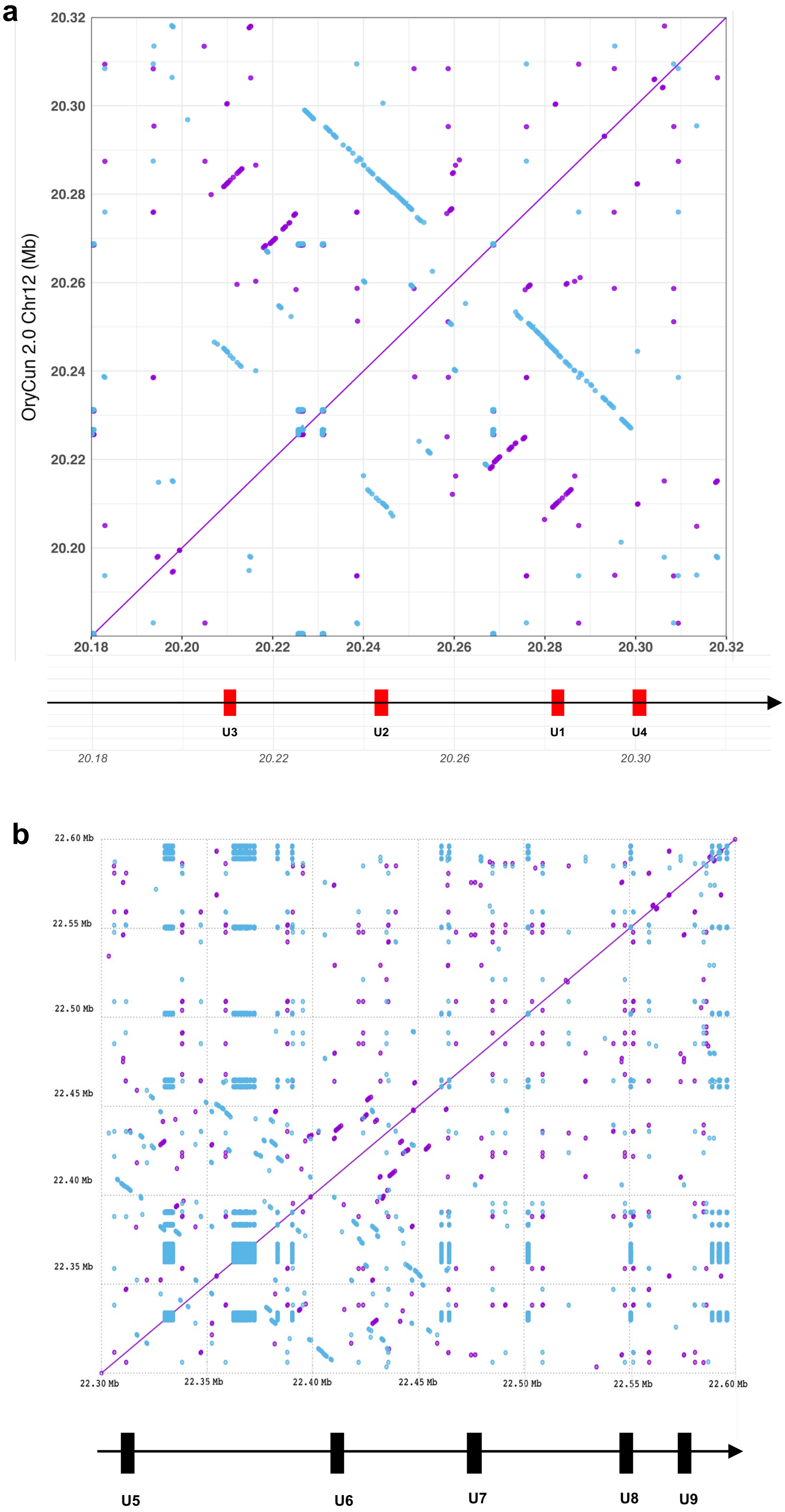


**Fig. S2. Duplications within the MHC-I region.** (**a**) The 0.14 Mb sequence of the first MHC-I region (chr12: 20,180,000-20,320,000) was compared to itself, and each dot marks an exact match of 50 bp or greater. **(b)** The 0.3 Mb sequence covering the second cluster of the MHC-I genes (chr12: 22,300,000-22,600,000) was compared to itself, and each dot marks an exact match of 50 bp or greater.

| | |
|---|---|
| M | Methionine start of SP |
| TWA/TQA/TRA/VGA | End of SP |
| GSH/ASH | Beginning of a1/a2 domain |
| Y/T/K/W | Invariant residues |
| N-X-S/T | Glycosylation motif |
| ED/AA/EA/DT | Loop |
| C/CHV | Conserved cysteine |
| DPP | Loop in a3 domain |
| WQRDG | Conserved Ig-like domain motif |
| TLRW | End of immunoglobulin domain |
| SS/PA/PS | Loop |
| IVG | Beginning of TM region |
| RRK/RRR/RKS | Charged residues |
| SS-X-X | Beginning of Cytoplasmic region |
| Y-X-X-Ø | Conserved tyrosine-based YXXØ motif |

```
Signal Peptide
HLA-A      -MAV---------MAPRTLLLLLSGALALTQTWA
Orcu-U1    -MGS---------MAPRTLLLLLAGALTLKDTQA
Orcu-U2    -------------MAPGTLLLLLAGALTLTQTRA
Orcu-U3    MMKS---------LAPRTLLLLLPGALVLTETRE
Orcu-U4    -------------MELRILLLMFCGALARPTTRA
Orcu-U5    -MFS---------LMPLTLLLLLAGALTLTQTQA
Orcu-U6    MMGS---------MAPGTLLPLLAGALTLTETWA
Orcu-U7    -MGT---------LGLRP-LLLLSGALVLAETRA
Orcu-U8    -------------MECGALLLLLAGALTLTETRT
Orcu-U9    --MAARAHRGPPTWAGPWALVFLL-LSFQRDVGA

a1 Domain            10        20        30        40        50        60        70        80                    90
HLA-A      GSHSMRYFFTSVSRPGRGEPRFIAVGYVDDTQFVRFDSDAASQRMEPRAPWIEQEGPEYWDQETRNVKAQSQTDRVDLGTLRGYYNQS------------EA
Orcu-U1    GSHSLRYFNIAVSRPGLGEPQYLEVGYVDDTQFVRFDSDAENPRMEPRAPWMRQVEPEYWDLNTRNAKSHEQTFRVNLNTLRGYYNQS------------AA
Orcu-U2    GLHSMSYFYTAVSRPGLGEPRFIAVGYVDDTQFVRFDSDAENPRAEQRAAWMRQVEPGYWDRNTQICKDTAQTFRVGLNTLRGYYNQS------------AA
Orcu-U3    GSHSMLYFETAVSRPGLGEPRFISVGYLDDTQFVRFDSDAENPREEPRAAWMRQVDPGYWDRNTQIYKDNAQTYRVDLNTLRGYYSQN------------AA
Orcu-U4    ASHSLRYFHVAVSRPGRGEAHYTSVGFVDDTQFVRFDSDAANPRVEPLAAWMDPEGPEYWDAQTKIAGVHAQTLRSNLRTALAYYNQS------------ED
Orcu-U5    GSHSMRFLHTTVSRAGLGEPRFTAVGYVDDTQCGRFDSDAASPRAEPREPWGRQVEPGFWDTQTQILKGEMLSYLEVLNSLRIYYNQS------------EA
Orcu-U6    GSHSMRHFFTSVSRPGLGEPRFLAVGYVDDTQFARFDSDAAAAGMEPRAPWMRQVEPRYWDMHRQRFEWGTLSFREYLSNLRIYYNQS------------EE
Orcu-U7    GSHSLQFFTTAVSRPGRGEARYITVGYVDDTPFARFDSYAENPRVEPRAPWMRQVEPGYWDRNTRFVKRSAQNYRVMLDNLRGYYNQS------------EA
Orcu-U8    GSHSLRYFTTAVSRPGLPEPRFMSVGYVDDTQFVRFDSDAASPRAEPRAPWMQQVEPWYWDRQTERAKNTALSFRVNLQTLRGYYNQS------------DT
Orcu-U9    GSHSLQYFYWAVTEPSPGVPAFAASGFVDNQRFIHYASEE--RKAKPCVQWLR-EEPSYFEDETKIFTSRMKIFQLSLRNVQQYYNQTLRPSSKPEQPHKLA

a2 Domain             100       110       120       130       140       150       160       170       180
HLA-A      GSHTIQIMYGCDVGSDGRFLRGYRQDAYDGKDYIALNEDLRSWTAADMAAQITKRKWEAAHEA-EQLRAYLDGTCVEWLRRYLENGKETLQRT
Orcu-U1    GSHTIQWMYGCEVGADGRLLRGYRQDAYDGADYIALNEDLRSWTAADTAAQNTRRKWEAAGAA-ERQRAYLEGECVEWLRRYLEMGKETLQRA
Orcu-U2    GSHTLQSMFGCEVWEDGRFFCGYTQYAYDGADYLALNEDLRSWTAADAAAQNTQRKWEAAGAA-ESHRAYLEGTCVEWLRRYLEMGKETLQRA
Orcu-U3    GSHTIQRMYGCEVGADGRFLRGYQQDAYDGADYIALNEDLRSWTAADTAAQITRRKWEAAGAA-EQDRAYLGGTCVEWLRRYLEMGKETLQRA
Orcu-U4    GSHSFQWTSGCDVGPDGRFLRGYEEFAYDGADYIVLNEDLRSWTAADMAAQITRRKWEALGHA-EHYRAYLQRECVEWLRRYLEMGKEALQRA
Orcu-U5    GSHTLQSMFGCEVGPDGRFLRGYSQYAYDGADYLVLNEHLHSWTVVDTRPNIWKPEWD----G-EQCTSFLEGECVEWLRGFLEMGKETLQRV
Orcu-U6    GSHTIQGMYGCEVGPDGRFLRGYSILAYDGADYLVLNEHLHSWTAVDTRPNIWKPEWD----G-KQCTSYLEGACVEWLRGFLEMGKETLQRV
Orcu-U7    GSHTYQSMCGCHLGPDGRLLRGYEQFAYDGADYITLNEDLRSWTAAHTAAQITQRQWEACGQA-ELRRSYLEGECVEWLRRYLEMGKETLQSP
Orcu-U8    GSHTLQWMYGCEVGADGRLLRGYEQFAYDGADYIALNEDLRSWTAADTAAQISKRKSEEASEA-ERHRSYLEGECVEWLRRYLQDGKETLQRA
Orcu-U9    GPHTLQFTYGCELQDDGGT-TGHWQYGYDGDDYLTLDVDSRQYVAASPLARYTKQKWEAGQDHMQRDVMYLTKECALWLHRYLELRGANSTRT

a3 Domain
HLA-A      DPPKTHMTHHPISDHEATLRCWALGFYPAEITLTWQRDGEDQTQDTELVETRPAGDGTFQKWAAVVVPSGEEQRYTCHVQHEGLPKPLTLRW
Orcu-U1    DPPKAHVTHHPASDREATLRCWALGFYPAEISLTWQRDGEDQTQDTELVETRPGGDGTFQKWAAVVVPSGEEQRYTCRVQHEGLPEPLTLTW
Orcu-U2    DPPKAHVTHHPASDKEATLRCWALGFYPAEISLTWQRDGEDQTQDTELVETRPGGDGTFQKWAAVVVPSGEEQRYTCRVQHEGLPEPLTLRW
Orcu-U3    DPPKAHVTHHPASDKEATLRCWALGFYPAEISLTWQRDGEDQTQDTELVETRPGGDGTFQKWAAVVVPSGEEQRYMCRVQHEGLPEPLTLRW
Orcu-U4    DPPKAHVTHHPASDRKATLRCWALGFYPAEISLTWQRDGEDQTQDTELVETRPGGDGTFQKWAAVVVPSGEEQRYTCHVQHEGLPEPLTLTW
Orcu-U5    DPPKTRVAHHPISDKEATLRCWALGFYPKEISLTWQRDGEDQTQDTELVETRPGGDGSFQKWAAVVVPSGEEQRYTCRVQHEGLPEPLTLRW
Orcu-U6    DPPKTHVAHHPISDKEATLRCWALGFYPKEISLTWQRDGEDQTQDTELVETRPGGDGSFQKWAAVVVPSGEEQRYTCRVQHEGLPEPLTLRW
Orcu-U7    VPPETRVTHHPISDQEATLRCWALGFYPKEISLTWQQDGEDQTQDTELVETRPGGDGTFQKWAAVVVPSGEEQRYTCRVQHEGLPEPLTLTW
Orcu-U8    DPPKAHVTHHPASDKEATLRCWALGFYPAEISLTWQRDGEDQTQDTELVETRPGGDGTFQKWAAVVVPSGEEQRYTCRVQHEGLPEPLTLRW
Orcu-U9    DPPKAHVTHHPASDKEATLRCWALGFYPAEISLTWQRDGEDQTQDTELVETRPGGDGTFQKWAAVVVPSGEEQRYTCRVQHEGLPEPLTLRW

Transmembrane and Cytoplasmic Domain
HLA-A      ELSSQPTIPIVGIIAGLVLLGAVITGAVVAAVMWRRKSSGGEGVKDRKGGSYTQAA---------SSDSAQGSDVSLTACKV
Orcu-U1    EPPAQPTALIVGIVAGVLGVL-LILGAVVAVVMRRKHSSDG------KGGKYAPAA---------GGHRDQGSDDSLMP
Orcu-U2    EPPAQPTALIVGIVAGVLGVL-LILGVVVAVVRRRKLSSDG------KGGRYTPAA---------GDHWNQGSEDSLMP
Orcu-U3    EPPAQSTVLIVGIVAGVLILGAMVPVVMRWMRTRRRQSSDG------RGGRYTPAAVKFWRVMLGGGHRDPCHSEKPMETLPAHPTLPPQPRPLWTSQSPDLFTPGGGIALSL
Orcu-U4    EPPSQSTNSIVI------TVCLVLLGAVIAFVLWWRKSSG
Orcu-U5    EPPAQPTTLSVGVVAGLVLG-AVVTAAVVAFVMRKRKSSVN------NGRRYAEST---------GSDSAQDSDMSLTASTA
Orcu-U6    EPPAQPTTLIVGVVAGLVLG-AVVTAAVVAFVMRKRKSSVN------NGRRYAEST---------DSKNSQDSHMSLTASTATA
Orcu-U7    ERSSQPTALIVGTIAGL-ILGAVVTAAMLIAVIKRKKSSGG------EGGSYTQAA---------SSDSTQDSDGSLTASTV
Orcu-U8    EPPAQPTALIVGIVAGVHGVL-LILGAVVAVVMRRKLSSDG------KGGKYAPAA---------GGHRDQGSDDSLLSVKP
Orcu-U9    EPPAQPTALIVGIVAGVLGVL-LILGVVVAVVRRRKHSSGS------KGWS
```

**Fig. S3. Annotated amino acid sequence alignment of nine rabbit MHC-I genes and human HLA-A.** Triangles mark the position of introns. Residue numbering in α1 and α2 domains is based on HLA-A. Conserved motifs and residues are coloured.

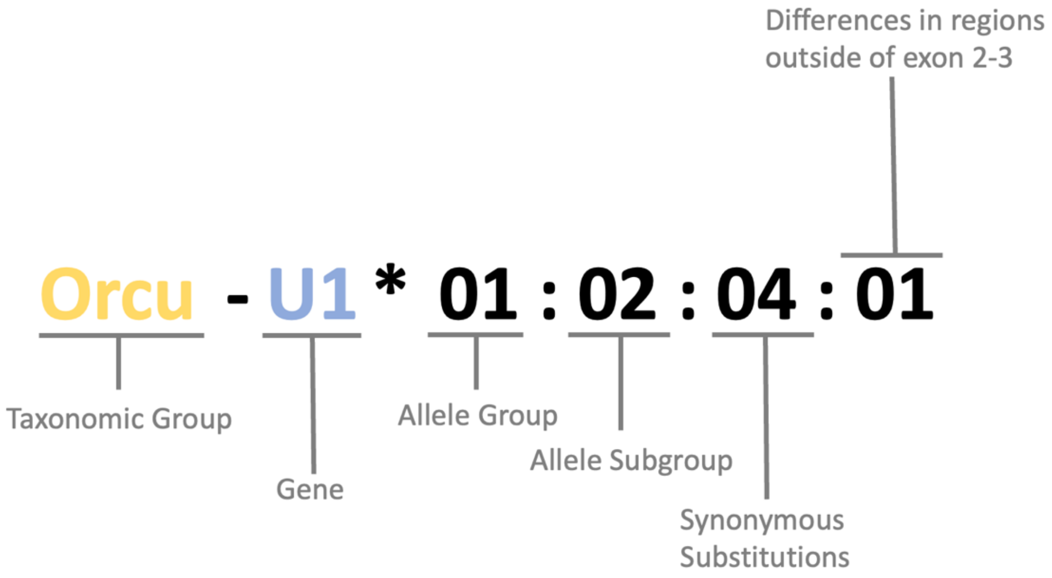


**Fig. S4. Schematics of the nomenclature of rabbit MHC alleles.** Yellow code represents the taxonomic group/species and blue indicates the name of the MHC-I gene, divided by a hyphen from the four-letter taxonomic code. The gene designation is followed by an asterisk, followed with an allelic string composed of four sets of digits, separated by colons. The four sets of digits indicate the allele group, allele subgroup, and synonymous substitutions, and nucleotide difference outside the peptide-binding region if information available.

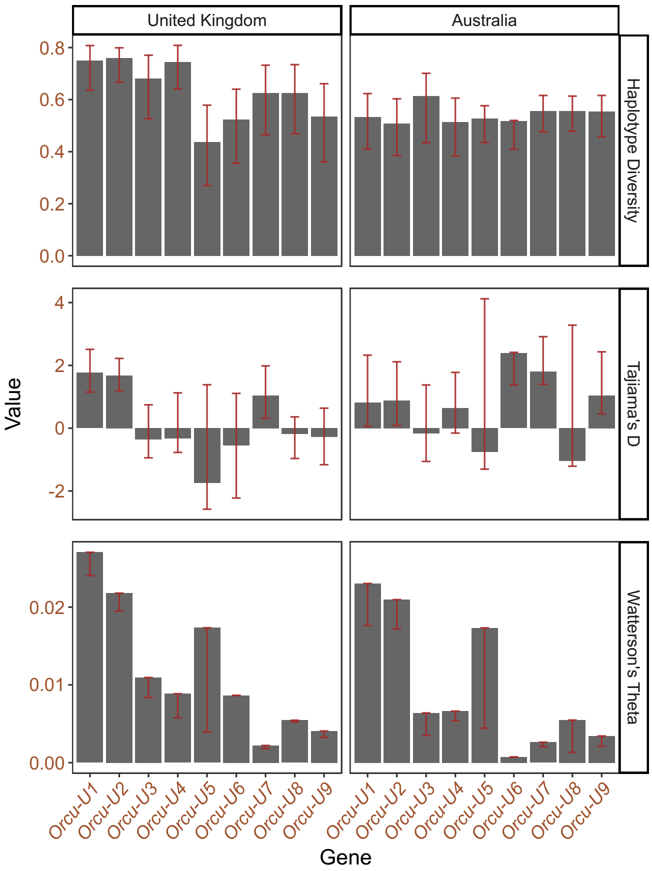


**Fig. S5. Haplotype diversity, Tajima's D and Watterson's Theta statistics of 9 MHC-I genes in the modern samples from Britain and Australia.** Confidence intervals correspond to the 0.025 and 0.975 quantiles of 1,000 bootstrap replicates estimates obtained by resampling alleles with replacement.

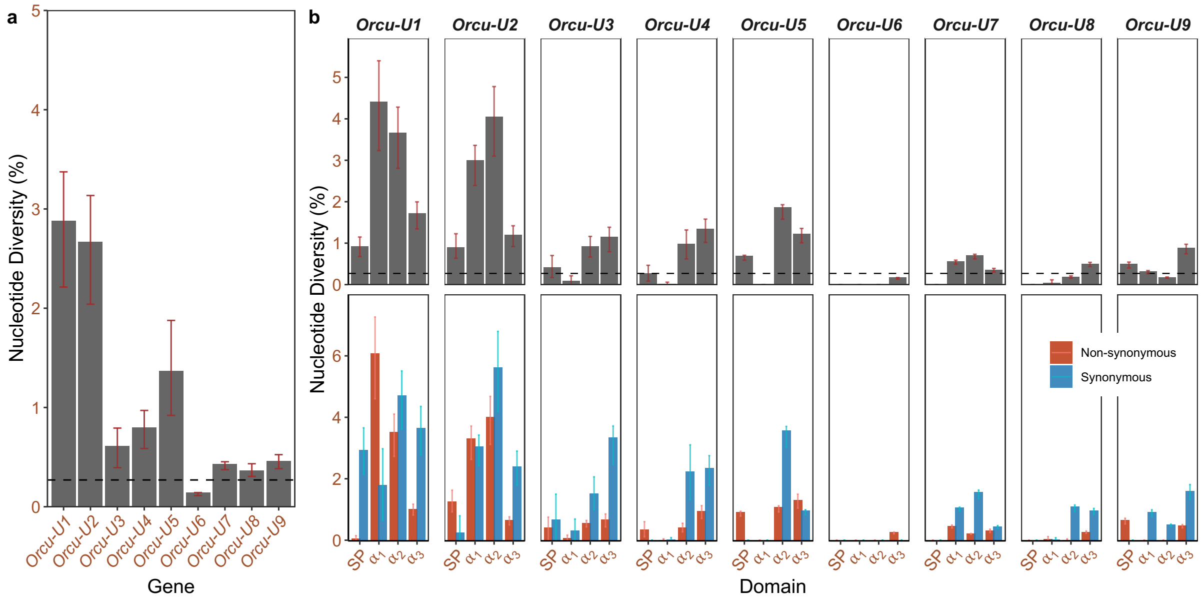


**Fig. S6. Genetic diversity of different MHC-I genes in Australia.** (**a**) Nucleotide diversity of 9 MHC-I genes in Australia. (**b**) Nucleotide diversity, non-synonymous and synonymous nucleotide diversity of different exons encoding different domains in Australia. The domains are the signal peptide (SP), the $\alpha_1$ and $\alpha_2$ domains which together form the peptide-binding groove, and the transmembrane-proximal $\alpha_3$ domain. The top panel shows total diversity, while the bottom panel distinguishes between non-synonymous and synonymous diversity. The dashed horizontal line on panels **(a)** and the top row of **(b)** indicates the mean genome-wide nucleotide diversity. Confidence intervals in **(a)** and **(b)** correspond to the 0.025 and 0.975 quantiles of 1,000 bootstrap replicates estimates obtained by resampling alleles with replacement.

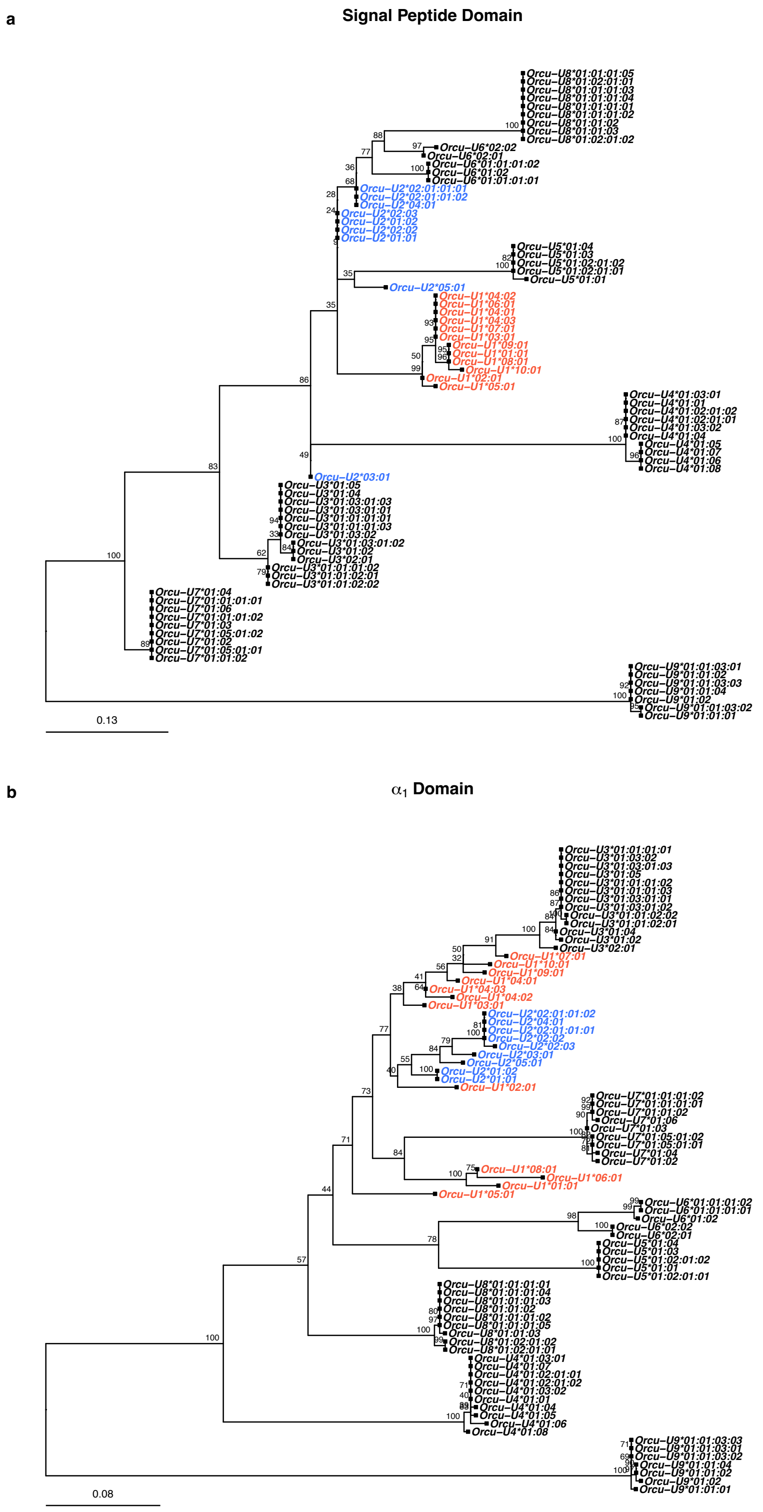
a
Signal Peptide Domain
Orcu−U8*01:01:01:05
Orcu−U8*01:02:01:01
Orcu−U8*01:01:01:03
Orcu−U8*01:01:01:04
Orcu−U8*01:01:01:01
Orcu−U8*01:01:01:02
Orcu−U8*01:01:02
Orcu−U8*01:01:03
Orcu−U8*01:02:01:02
Orcu−U6*02:02
Orcu−U6*02:01
Orcu−U6*01:01:01:02
Orcu−U6*01:02
Orcu−U6*01:01:01:01
Orcu−U2*02:01:01:01
Orcu−U2*02:01:01:02
Orcu−U2*04:01
Orcu−U2*02:03
Orcu−U2*01:02
Orcu−U2*02:02
Orcu−U2*01:01
Orcu−U5*01:04
Orcu−U5*01:03
Orcu−U5*01:02:01:02
Orcu−U5*01:02:01:01
Orcu−U5*01:01
Orcu−U2*05:01
Orcu−U1*04:02
Orcu−U1*06:01
Orcu−U1*04:01
Orcu−U1*04:03
Orcu−U1*07:01
Orcu−U1*03:01
Orcu−U1*09:01
Orcu−U1*01:01
Orcu−U1*08:01
Orcu−U1*10:01
Orcu−U1*02:01
Orcu−U1*05:01
Orcu−U4*01:03:01
Orcu−U4*01:01
Orcu−U4*01:02:01:02
Orcu−U4*01:02:01:01
Orcu−U4*01:03:02
Orcu−U4*01:04
Orcu−U4*01:05
Orcu−U4*01:07
Orcu−U4*01:06
Orcu−U4*01:08
Orcu−U2*03:01
Orcu−U3*01:05
Orcu−U3*01:04
Orcu−U3*01:03:01:03
Orcu−U3*01:03:01:01
Orcu−U3*01:01:01:01
Orcu−U3*01:01:01:03
Orcu−U3*01:03:02
Orcu−U3*01:03:01:02
Orcu−U3*01:02
Orcu−U3*02:01
Orcu−U3*01:01:01:02
Orcu−U3*01:01:02:01
Orcu−U3*01:01:02:02
Orcu−U7*01:04
Orcu−U7*01:01:01:01
Orcu−U7*01:06
Orcu−U7*01:01:01:02
Orcu−U7*01:03
Orcu−U7*01:05:01:02
Orcu−U7*01:02
Orcu−U7*01:05:01:01
Orcu−U7*01:01:02
Orcu−U9*01:01:03:01
Orcu−U9*01:01:02
Orcu−U9*01:01:03:03
Orcu−U9*01:01:04
Orcu−U9*01:02
Orcu−U9*01:01:03:02
Orcu−U9*01:01:01
0.13
b
α1 Domain
Orcu−U3*01:01:01:01
Orcu−U3*01:03:02
Orcu−U3*01:03:01:03
Orcu−U3*01:05
Orcu−U3*01:01:01:02
Orcu−U3*01:01:01:03
Orcu−U3*01:03:01:01
Orcu−U3*01:03:01:02
Orcu−U3*01:01:02:02
Orcu−U3*01:01:02:01
Orcu−U3*01:04
Orcu−U3*01:02
Orcu−U3*02:01
Orcu−U1*07:01
Orcu−U1*10:01
Orcu−U1*09:01
Orcu−U1*04:01
Orcu−U1*04:03
Orcu−U1*04:02
Orcu−U1*03:01
Orcu−U2*02:01:01:02
Orcu−U2*04:01
Orcu−U2*02:01:01:01
Orcu−U2*02:02
Orcu−U2*02:03
Orcu−U2*03:01
Orcu−U2*05:01
Orcu−U2*01:02
Orcu−U2*01:01
Orcu−U1*02:01
Orcu−U7*01:01:01:02
Orcu−U7*01:01:01:01
Orcu−U7*01:01:02
Orcu−U7*01:06
Orcu−U7*01:03
Orcu−U7*01:05:01:02
Orcu−U7*01:05:01:01
Orcu−U7*01:04
Orcu−U7*01:02
Orcu−U1*08:01
Orcu−U1*06:01
Orcu−U1*01:01
Orcu−U1*05:01
Orcu−U6*01:01:01:02
Orcu−U6*01:01:01:01
Orcu−U6*01:02
Orcu−U6*02:02
Orcu−U6*02:01
Orcu−U5*01:04
Orcu−U5*01:03
Orcu−U5*01:02:01:02
Orcu−U5*01:01
Orcu−U5*01:02:01:01
Orcu−U8*01:01:01:01
Orcu−U8*01:01:01:04
Orcu−U8*01:01:01:03
Orcu−U8*01:01:02
Orcu−U8*01:01:01:02
Orcu−U8*01:01:01:05
Orcu−U8*01:01:03
Orcu−U8*01:02:01:02
Orcu−U8*01:02:01:01
Orcu−U4*01:03:01
Orcu−U4*01:07
Orcu−U4*01:02:01:01
Orcu−U4*01:02:01:02
Orcu−U4*01:03:02
Orcu−U4*01:01
Orcu−U4*01:04
Orcu−U4*01:05
Orcu−U4*01:06
Orcu−U4*01:08
Orcu−U9*01:01:03:03
Orcu−U9*01:01:03:01
Orcu−U9*01:01:03:02
Orcu−U9*01:01:04
Orcu−U9*01:01:02
Orcu−U9*01:02
Orcu−U9*01:01:01
0.08

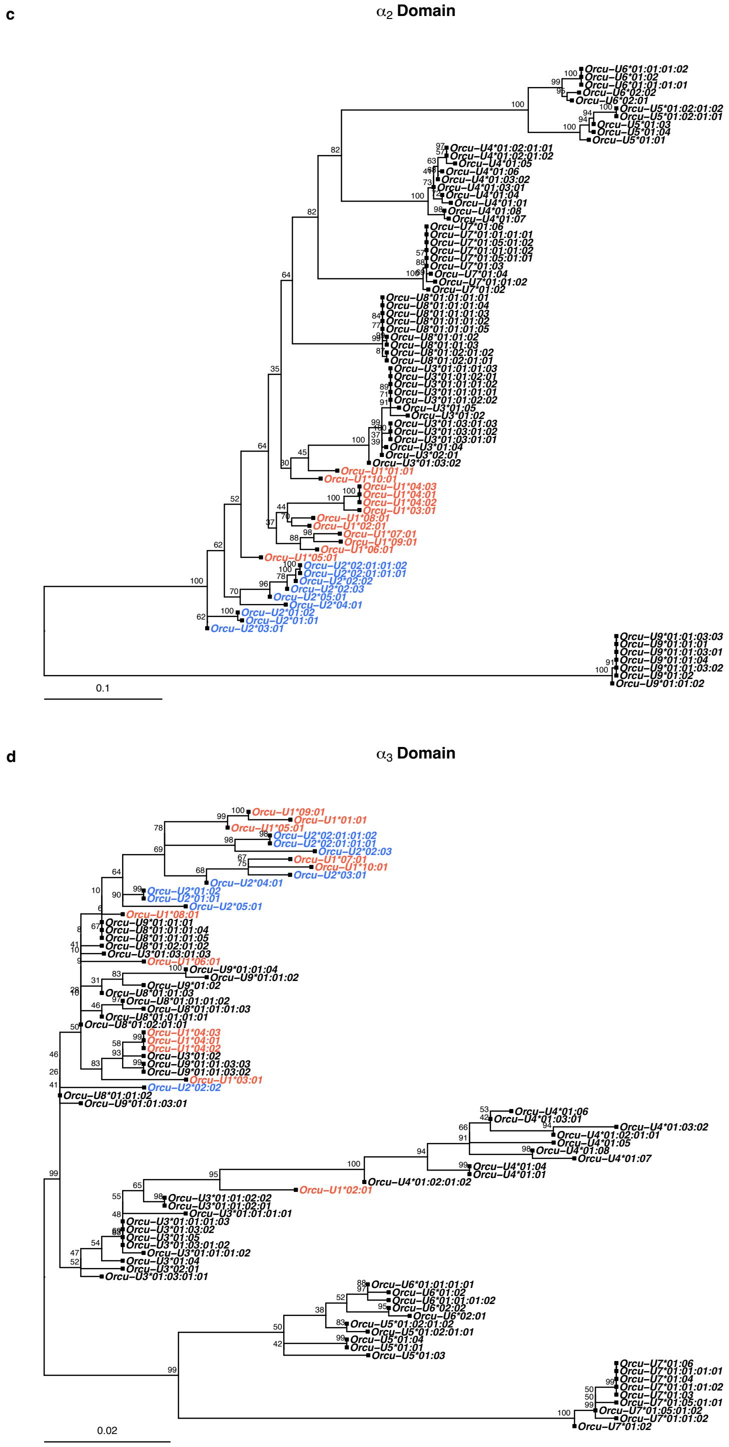


**Fig. S7. Phylogenetic analyses of different domains from 9 MHC-I candidates.** Phylogeny of nucleotide sequences of the signal peptide, α1, α2 and α3 domains (in (**a**)-(**d**), respectively) from 9 MHC-I genes constructed by maximum likelihood using JC69 model. Branch support is indicated as percentage of trees out of 1,000 bootstrap replicates. The alleles coloured red and blue stand for *Orcu-U1* and *Orcu-U2*.

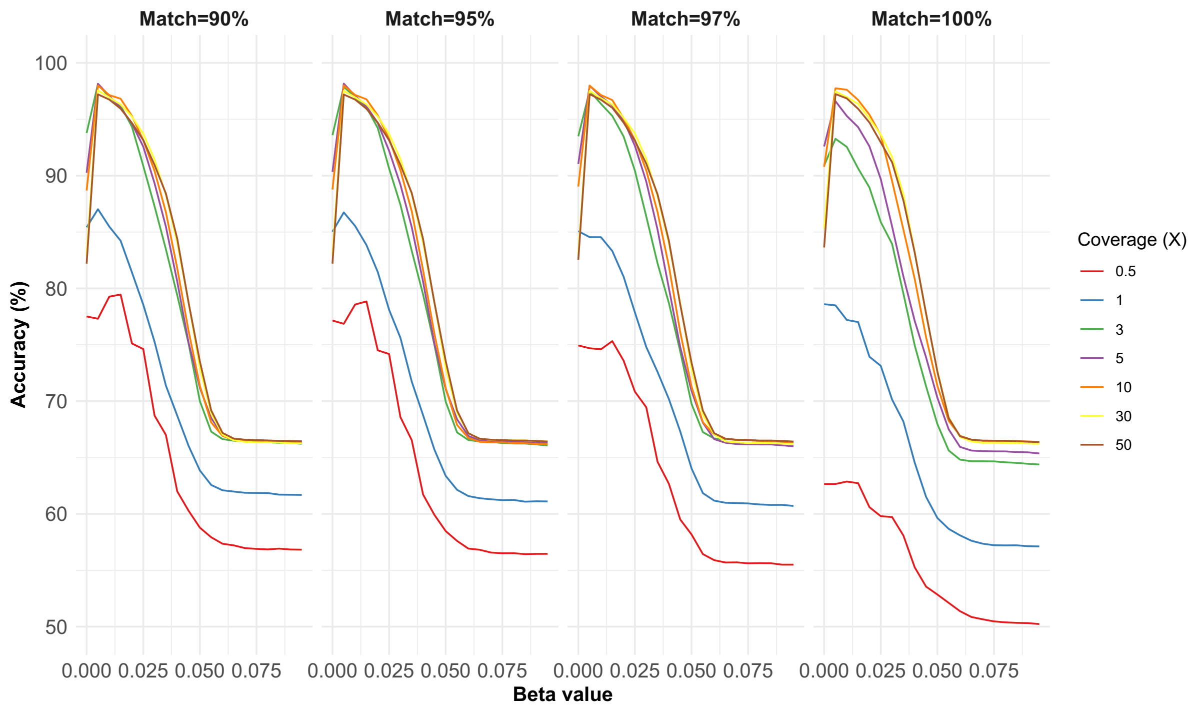


**Fig. S8. Accuracy of genotyping MHC-I from historical DNA sequencing.** Sequence reads were simulated to match the patterns of degradation and post-mortem damage observed in our historical dataset, and to match the genotypes of *Orcu-U1* to *Orcu-U8* observed in our modern samples used for the PacBio transcript and amplicon sequencing. Genotypes were then called by modifying the method of [33] for rabbits. Facets represent the resulting accuracy for different matching tolerance of mapping: 90%, 95%, 97% and 100%. Beta is a parameter that penalises the calling of heterozygotes to avoid heterozygosity bias. The accuracy is the percentage of *Orcu-U1* and *Orcu-U2* alleles correctly called across all the simulated samples.

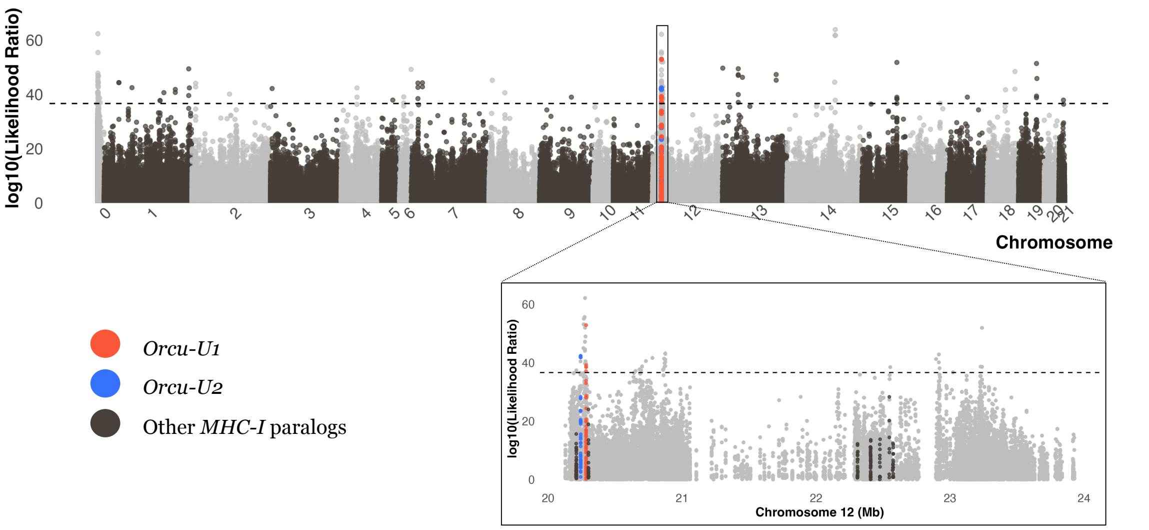


**Fig. S9. Genome-wide selection scan based on allele frequency changes after the introduction of MYXV. (a)** Selection scan assuming selection was the same across three populations. The upper side plots show the genome-wide scale of selection pattern. The bottom-side plots are the zoomed in observations on MHC-I region, with the red and blue dots showing variants within the exon region of *Orcu-U1* and *Orcu-U2*, and the brown dots representing the variants within the exon regions of the other seven MHC-I paralogs. In all plots, the dashed black lines indicate the genome-wide 95% significance threshold from permuting sample collection dates within each country 1,000 times.

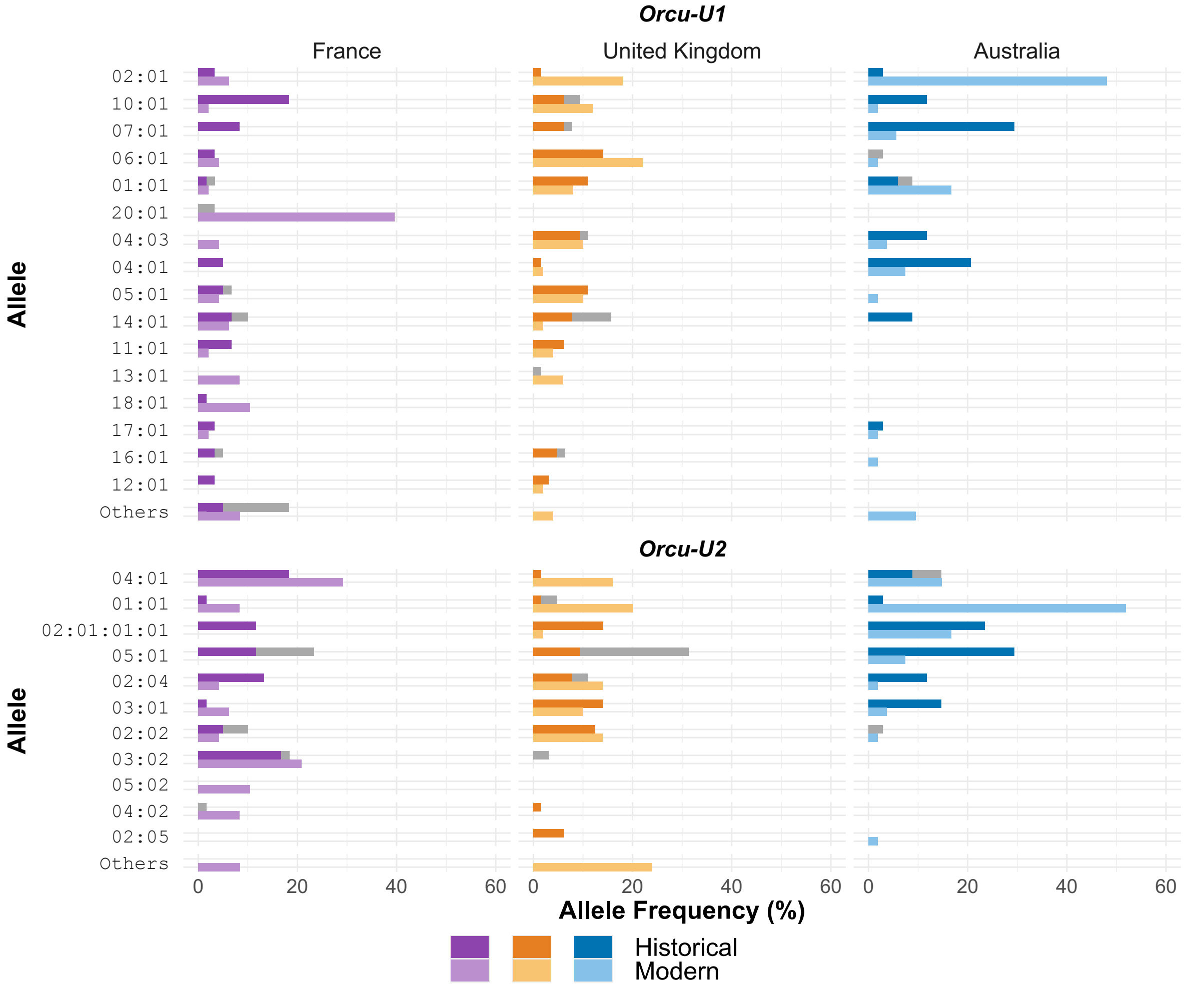


**Fig. S10. Allele frequency changes before and after myxomatosis within three populations.** The allele frequencies for each allele in gene *Orcu-U1* and *Orcu-U2*, separating in three studied populations France, Australia, and Britain. Alleles ordered by overall frequency. The darker bar within each facet stands for the frequencies of historical individuals while the lighter colour indicates for the modern ones. The unsatisfactory calls of alleles from historical specimens are represented with stacked grey bars (sequences that resemble but are not identical to that allele in our database). Alleles with a combined frequency of less than 8% across all categories are grouped together under 'Others'.

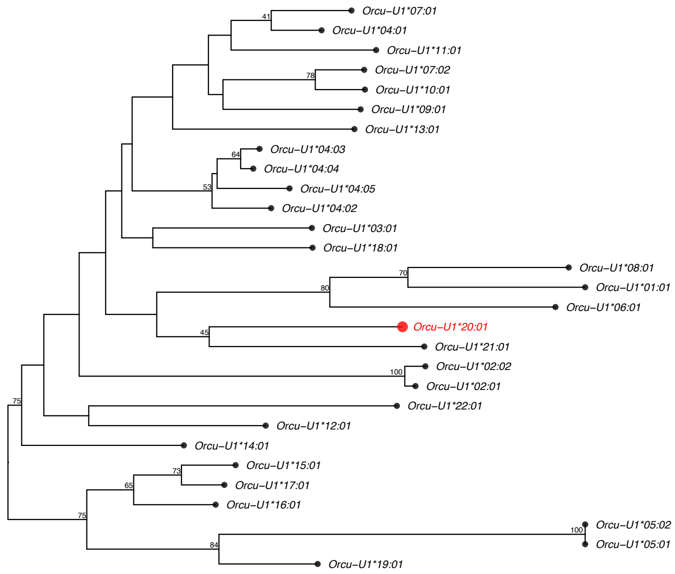


**Fig. S11. Neighbour-joining phylogenetic tree of 29 *Orcu-U1* alleles constructed using the JTT amino acid substitution model.** Bootstrap support values (1000 replicates) ≥40% are displayed at internal nodes. The allele *Orcu-U1**20:01, which showed population-specific increase in the modern French population, is highlighted in red.

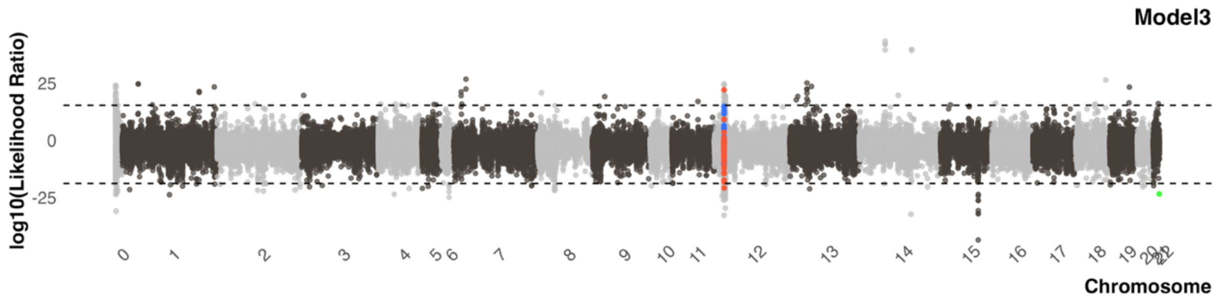


**Fig. S12. Population-specific selection scan for the MHC-I haplotype combining *Orcu-U1**02:01 and *Orcu-U2**01:01.** The plot shows the likelihood ratio statistic for the combined haplotype (green dot) located on chromosome 22 for visualisation purposes, compared to genome-wide variants. Positive values and negative values show sites selection acted on in all three populations or just one population, respectively. Red and blue dots showing variants within the exon region of *Orcu-U1* and *Orcu-U2*, and the dashed black lines indicate the genome-wide 95% significance threshold from permuting sample collection dates within each country 1,000 times.

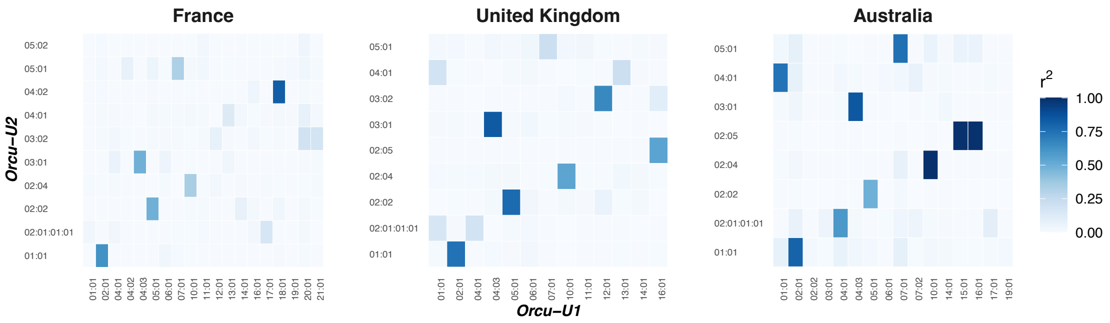


**Fig. S13. Pairwise linkage disequilibrium between alleles of *Orcu-U1* and *Orcu-U2*.** The heatmap shows the pairwise linkage disequilibrium of alleles from *Orcu-U1* and *Orcu-U2* with frequency over 0.01*,* assessed using $r^2$ values, with the darker blue showing a LD close to 1.

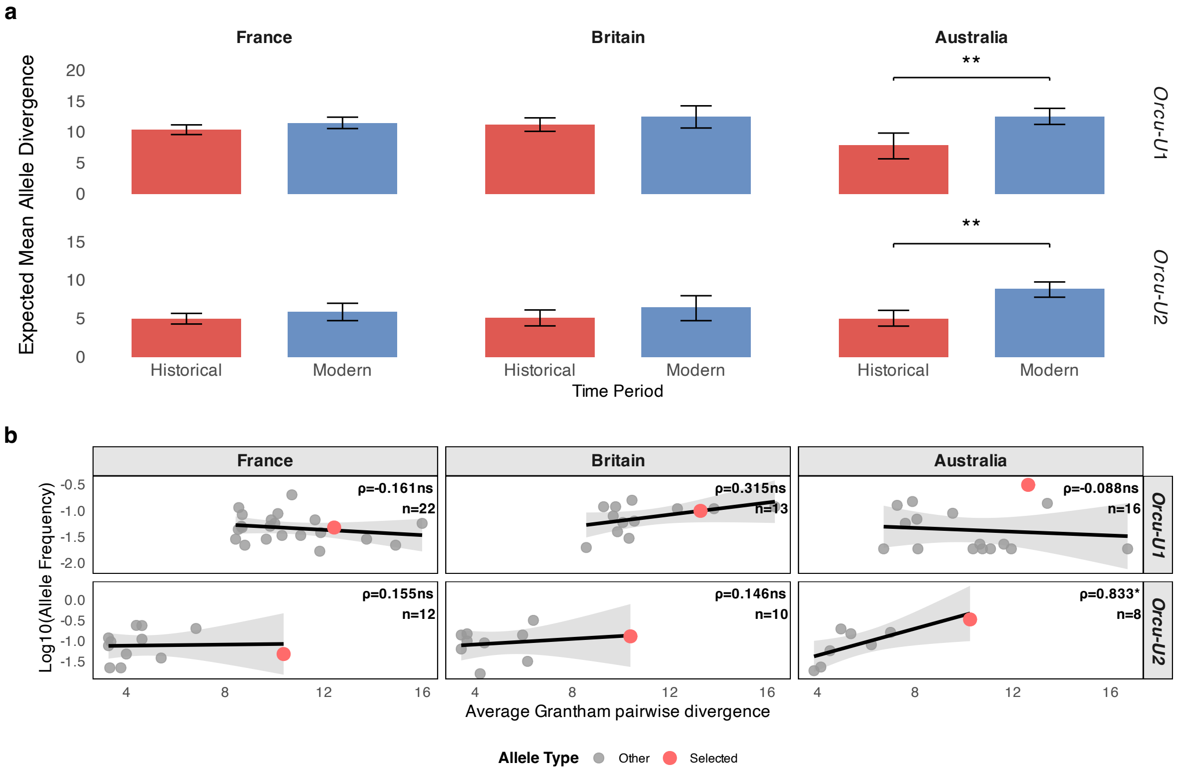


**Fig. S14. Changes in allele divergence across populations and time periods.** (**a**) Bars represent expected mean allele divergence of *Orcu-U1* and *Orcu-U2* in historical and modern samples calculated as the Hardy-Weinberg weighted average of pairwise Grantham distances between all alleles. Error bars show 95% confidence intervals estimated via 1,000 bootstrap replicates by resampling samples with replacement. Statistical significance of differences between historical and modern groups was assessed using 1,000 permutation tests where time period labels were randomly reassigned to individuals to generate a null distribution (**p<0.01). (**b**) Correlation between average Grantham pairwise divergence to common alleles (≥5% frequency) and allele frequency for *Orcu-U1* and *Orcu-U2* across France, UK, and Australia populations. Selected alleles *(Orcu-U1*02:01, Orcu-U2**01:01) are highlighted in red. Spearman correlation coefficients ($\rho$) are shown for each gene-population combination (*$p$<0.05; *ns* = not significant).

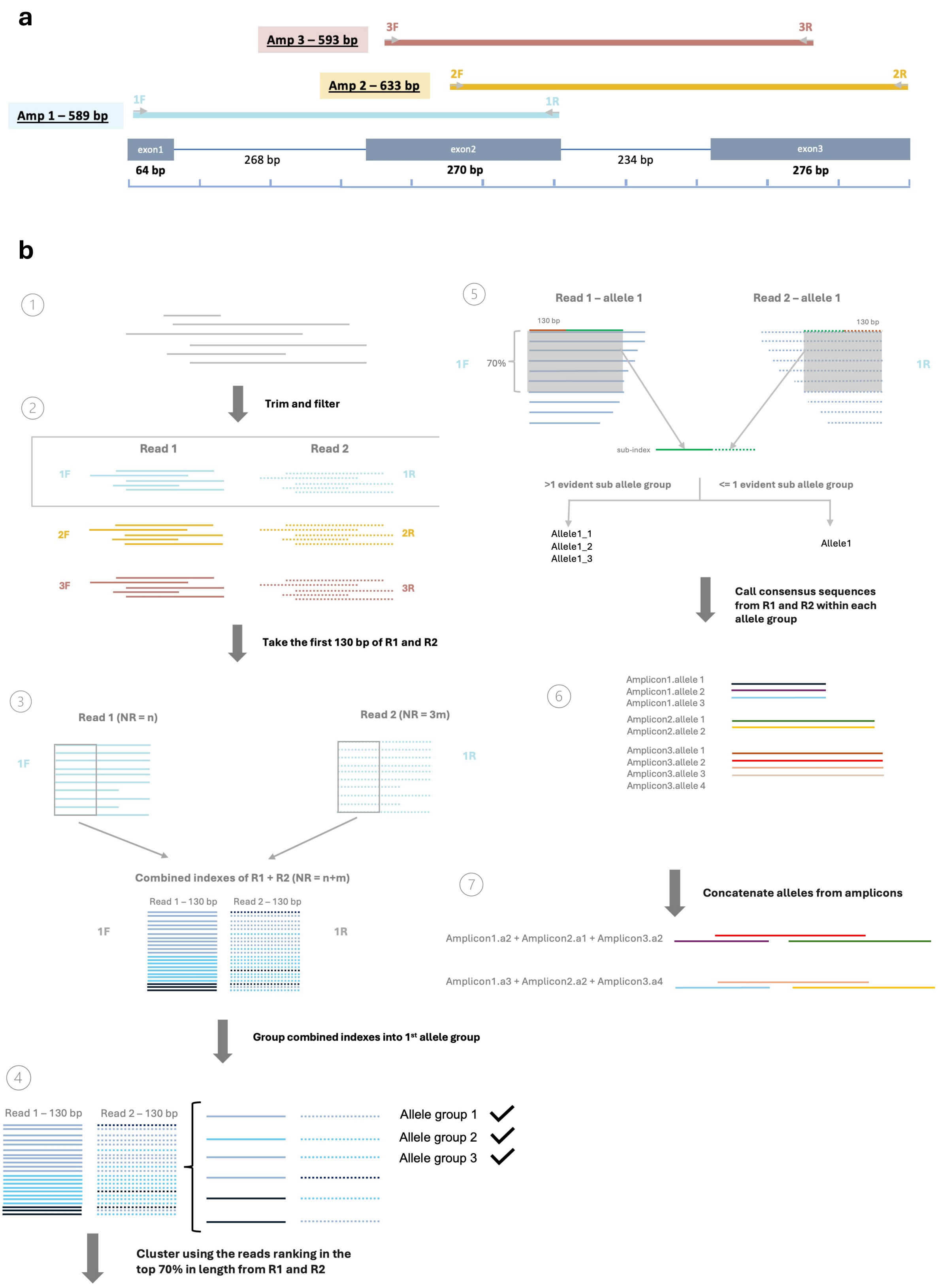


**Fig. S15. Design of MiSeq amplicon sequencing pipeline.** (**a**) Design of the MiSeq amplicon sequencing capturing the exon1-3 region of both *Orcu-U1* and *Orcu-U2*. The design includes three amplicons, with each two of them having overlapped regions. (**b**) Bioinformatics pipeline of the MHC-I allele typing from amplicon sequencing data.

| Gene Name | Associated *Ensembl* gene IDs[1] | *Ensembl* transcript IDs[2] | Transcriptional orientation | Number of samples | Alteration of annotations[3] | Number of alleles |
|---|---|---|---|---|---|---|
| *Orcu-U1* | ENSOCUG00000010972 | ENSOCUT00000033804 | - | 234 | N | 29 |
| *Orcu-U2* | ENSOCUG00000011016 | - | + | 234 | Y | 17 |
| *Orcu-U3* | ENSOCUG00000010972 | ENSOCUT00000043824 | - | 68 | N | 13 |
| *Orcu-U4* | ENSOCUG00000013304 | - | - | 68 | Y | 10 |
| *Orcu-U5* | ENSOCUG00000006557 | - | + | 68 | Y | 5 |
| *Orcu-U6* | ENSOCUG00000010984 | - | - | 68 | Y | 5 |
| *Orcu-U7* | - | - | + | 68 | Y | 9 |
| *Orcu-U8* | ENSOCUG00000008993 | - | + | 68 | Y | 9 |
| *Orcu-U9* | - | - | + | 68 | Y | 7 |

**Table S1. Annotation of rabbit MHC-I genes.**[1] The gene annotation in the *Ensembl* database that includes some or all the gene. [2] The transcript ID that corresponds to the manually annotated genes. Other transcripts were not confirmed. [3] Where none of the transcripts are correct and the gene was manually reannotated. The gene underlined were included in the probe design for PacBio transcriptome sequencing.

| Tissue | SRR ID | BioProject Acc. |
|---|---|---|
| Aorta | SRR1786004 | PRJNA274427 |
| Aorta | SRR1786005 | PRJNA274427 |
| Aorta | SRR1786006 | PRJNA274427 |
| Blood | SRR388309 | PRJNA78323 |
| Brain | SRR388305 | PRJNA78323 |
| Embryo | SRR1789369 | PRJNA274427 |
| Embryo | SRR1789370 | PRJNA274427 |
| Embryo | SRR1789384 | PRJNA274427 |
| Heart | SRR1789085 | PRJNA274427 |
| Heart | SRR1789162 | PRJNA274427 |
| Heart | SRR1789374 | PRJNA274427 |
| Kidney | SRR32117905 | PRJNA1027777 |
| Kidney | SRR32117904 | PRJNA1027777 |
| Liver | SRR1789057 | PRJNA274427 |
| Liver | SRR1789061 | PRJNA274427 |
| Liver | SRR388316 | PRJNA78323 |
| Lung | SRR388308 | PRJNA78323 |
| Ovary | SRR388314 | PRJNA78323 |
| Skeletal Muscle | SRR388304 | PRJNA78323 |
| Skeletal Muscle | SRR32117903 | PRJNA1027777 |
| Skin | SRR388313 | PRJNA78323 |
| Spleen | SRR32117902 | PRJNA1027777 |
| Spleen | SRR32117901 | PRJNA1027777 |
| Spleen | SRR32117900 | PRJNA1027777 |
| Testis | SRR388295 | PRJNA78323 |

**Table S2. Samples used for Tissue-specific expression analysis of MHC-I genes.** The NCBI SRR accession numbers for all the samples included in the tissue-specific expression analysis. The samples with their IDs underlined were generated in our lab for this study.

| Name | Sequence |
|---|---|
| 1F | TCGTCGGCAGCGTCAGATGTGTATAAGAGACAGGGACCCTCCTCYTGCTGCTC |
| 1R | GTCTCGTGGGCTCGGAGATGTGTATAAGAGACAGCCGCGCTCTGGTTGTAGTAG |
| 2F | TCGTCGGCAGCGTCAGATGTGTATAAGAGACAGCGTGGATGSGGCAGGTGG |
| 2R | GTCTCGTGGGCTCGGAGATGTGTATAAGAGACAGCTGCGCGCTGCAGYGTCT |
| 3F | TCGTCGGCAGCGTCAGATGTGTATAAGAGACAGCGTGTCCCGGCCCGGCCTGGG |
| 3R | GTCTCGTGGGCTCGGAGATGTGTATAAGAGACAGGCAGGTCCTCGTTCAGGGCG |

**Table S3. Design of the MiSeq amplicon sequencing capturing the exon1-3 region of both *Orcu-U1* and *Orcu-U2*.** Each sequence consists of Illumina adaptor followed by a gene-specific primer (underlined).

**Supplementary Files**

Data file S1 – List of all samples used in this study and sequencing metrics for exome sequencing data.

Data file S2 – List containing the coordinates of exons of nine annotated MHC-I genes.

Data file S3 – Genotyping results of all individuals included in this study.

Data file S4 – Sequence alignment used for generating phylogeny.

Data file S5 – List containing nomenclature of identified MHC-I alleles from 9 genes.

Data file S6 – Phased haplotypes their frequencies at *Orcu-U1* and *Orcu-U2* loci.

Data file S7 – Likelihood ratios and parameter estimation for haplotype trajectories over time.

Data file S8 – List containing PolydT primers of PacBio sequencing.

Data file S9 – BED file with probe coordinates for the exome capture.

Data file S10 – List containing the coordinates of genes within rabbit MHC region.
